# Fe-S protein assembly involves bipartite client binding and conformational flexibility in the CIA targeting complex

**DOI:** 10.1101/2020.02.01.930206

**Authors:** Susanne A. Kassube, Nicolas H. Thomä

## Abstract

The cytosolic iron sulfur (Fe-S) assembly (CIA) pathway is required for the insertion of Fe-S clusters into cytosolic and nuclear client proteins, including many DNA metabolism proteins. The molecular mechanisms of client protein recognition and Fe-S cluster transfer remain unknown. Here we report crystal structures of the CIA targeting complex and cryo-EM reconstructions of the complex bound either to the DNA replication factor primase or the DNA helicase DNA2. The structures, combined with biochemical, biophysical and yeast complementation assays, reveal an evolutionarily conserved, bipartite client binding mode facilitated by the structural flexibility of the MMS19 subunit. The primase Fe-S cluster is located ∼70 Å away from the catalytic cysteine in the CIA targeting complex, pointing to a conformationally dynamic mechanism of Fe-S cluster transfer. Altogether, our studies suggest a model for Fe-S cluster insertion and thus provide a mechanistic framework to understand the biogenesis of critical DNA replication and repair factors.

## Main

Iron-sulfur (Fe-S) clusters, found in all three domains of life^1^, stabilize protein folds, facilitate electron transfer processes, and sense oxygen or iron levels in cells. Many eukaryotic DNA replication and repair proteins, including B-family DNA polymerases α, δ, ε and ζ, the DNA primase large subunit (PriL), and DNA helicases XPD, RTEL1, FANCJ, DNA2, and DDX11, contain Fe-S clusters^2–8^. Although the precise roles of the Fe-S clusters in many of these proteins remain enigmatic, loss of the Fe-S cluster often leads to reduced protein stability and functionality^7, 9, 10^, and disease-causing mutations have been mapped to the Fe-S domains of XPD, FANCJ, and MUTYH^11^.

The biogenesis of eukaryotic Fe-S proteins is a highly regulated, multi-step process, which is conserved from yeast to human^12, 13^. The mitochondrial iron sulfur cluster (ISC) machinery is required for the biogenesis of all mitochondrial Fe-S proteins, while the cytosolic iron sulfur assembly (CIA) pathway is essential for the maturation of cytosolic and nuclear Fe-S proteins. The CIA pathway depends on the mitochondrial ISC machinery, which is thought to generate a sulfur-containing precursor that is exported into the cytosol^14^. As a first step in the CIA pathway, the [4Fe-4S] cluster is assembled on the NUBP1/NUBP2 scaffold complex (Nbp35-Cfd1 in yeast)^15–18^. The cluster is then thought to be transferred onto the intermediate carrier protein CIAO3 (Nar1 in yeast)^19, 20^. In the next step, the CIA targeting complex (CTC), consisting of MMS19, CIAO1, and CIAO2B (MET18, CIA1, and CIA2B in yeast; Fig. 1a), recognizes client apo-proteins through direct physical interactions and mediates the insertion of the Fe-S cluster. MMS19 is an adaptor subunit required for the recognition of diverse client proteins, including essential DNA replication and repair factors, by direct physical interactions^21^. Cells lacking MMS19 exhibit pleiotropic phenotypes including genome instability due to reduced levels of Fe-S cluster incorporation into DNA metabolism proteins and their consequent destabilization^22, 23^.

**Fig. 1.**
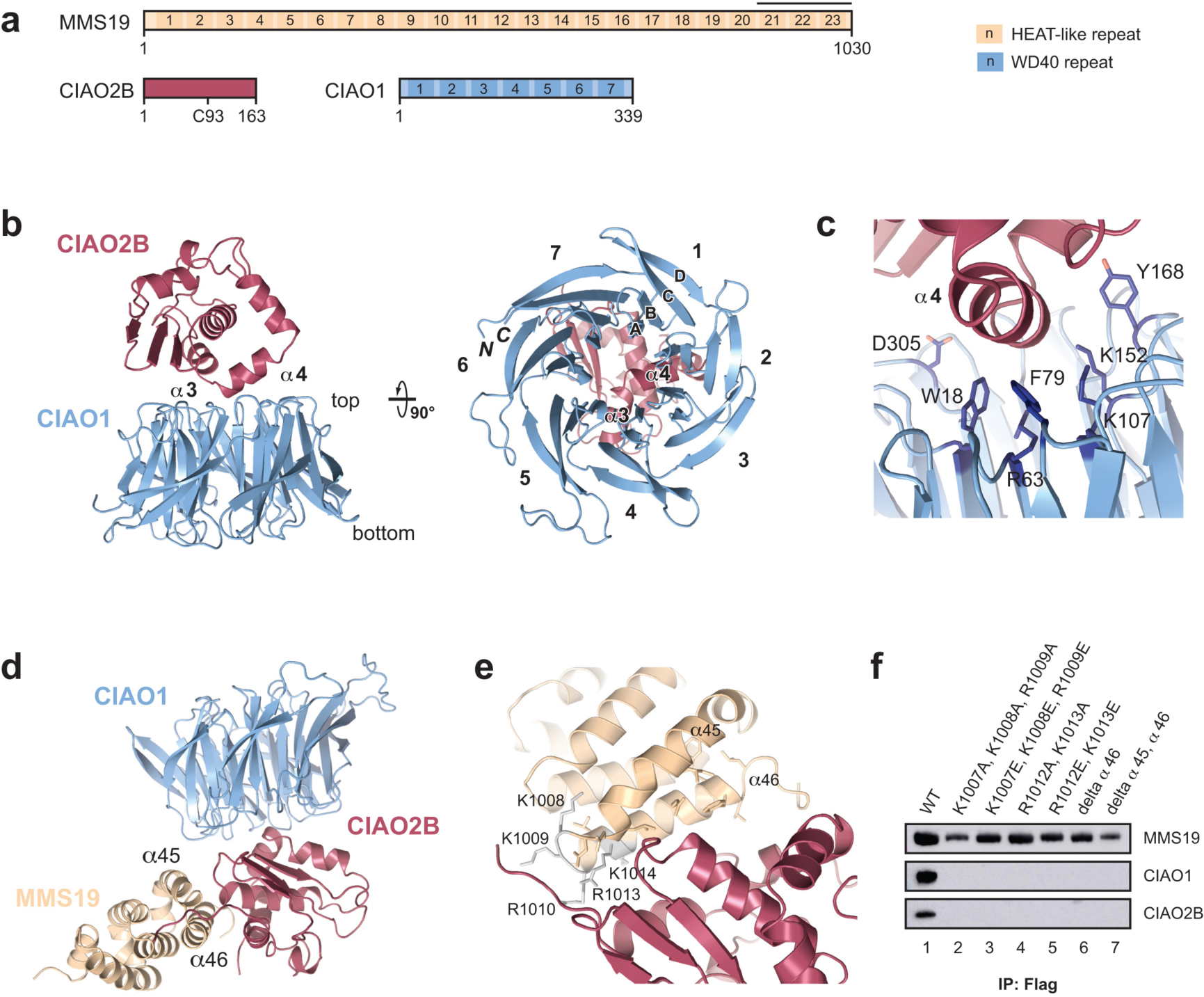
**Crystal structures of CIAO1-CIAO2B and MMS19^CTD^-CIAO1-CIAO2B complexes. a**, Domain organization of MMS19, CIAO2B, and CIAO1. The MMS19 fragment contained in the crystal structure in d is indicated with a bar above the last three HEAT-like repeats. **b**, Crystal structure of the *Drosophila* CIAO1-CIAO2B complex; *left* side view *right* view from the bottom face of the CIAO1 β-propeller; sheets A through D are labeled in blade 1. **c**, Close-up view of the interface showing CIAO1 amino acid side chains required for interaction, and CIAO2B helix α4. **d**, Crystal structure of the MMS19^CTD^-CIAO1-CIAO2B complex. **e**, Close-up view of the interface between MMS19^CTD^ and CIAO2B. **f**, Pull-down assay from HEK293 cells using Flag-tagged human MMS19 wt and mutants, and myc-tagged human CIAO2B/CIAO1.

CIAO2B, the smallest (17.6 kDa) CTC subunit, was previously shown to contain a highly reactive cysteine residue (Cys93 in human and Cys161 in yeast), and is therefore assumed to be the catalytically active component of the CTC^24^. Although structures of the CIAO2B-homolog FAM96A have been determined^25^, it remains unclear how exactly CIAO2B might facilitate the transfer of a preassembled Fe-S cluster. CIAO1 contains a seven-bladed WD40-repeat β-propeller domain^26, 27^, while MMS19, for which there is no structural information currently available, is predicted to be a large alpha-helical HEAT repeat protein^10, 28^.

Despite its essential cellular function, the overall architecture of the CTC, the specific functions of its subunits, and the molecular mechanism of Fe-S cluster transfer remain unclear. Given the structural and functional diversity of Fe-S proteins, and the lack of common sequence motifs or folds for Fe-S cluster coordination, the mechanisms by which the CTC specifically recognizes client proteins and assists in their maturation remain enigmatic. Our integrative structural and biochemical study provides detailed insights into the evolutionarily conserved mechanism of cytosolic and nuclear Fe-S protein biogenesis, and suggests a model for the transfer of Fe-S clusters from the intermediate carrier CIAO3 via the CTC and to diverse client proteins.

## Results

### Structure of CIAO1-CIAO2B catalytic core

While most Fe-S proteins involved in DNA replication and repair require the CTC adaptor protein MMS19 for their maturation, a subset of client proteins requires only the minimal hetero-dimeric CIAO1-CIAO2B “core” complex^10, 28^. We determined the structure of the *Drosophila melanogaster* minimal core complex by X-ray crystallography at a resolution of 2.0 Å (Fig. 1b, Extended Data Fig. 1 and Table 1). The structure reveals that CIAO2B interacts with the narrow-end face of the CIAO1 β-propeller, through contacts involving CIAO2B helices α3 and α4, with the N-terminal end of helix α4 positioned above the central cavity of the CIAO1 β-propeller. The CIAO2B interaction site, which is mostly formed by CIAO1 β-propeller blades 2, 3, and 4, is highly conserved from yeast to human (Extended Data Fig. 1b-e and Fig. 1c). To verify the interaction, we systematically mutated residues on the narrow-end face of CIAO1 and performed pull-down assays using insect cells over-expressing Strep-tagged CIAO2B and His-tagged CIAO1 (Extended Data Fig. 1d). All residues involved in the interaction with CIAO2B are located in the evolutionarily conserved patch formed by blades 2, 3, and 4 of CIAO1 and interact with helix α4 of CIAO2B (Fig. 1c).

**Table 1.**
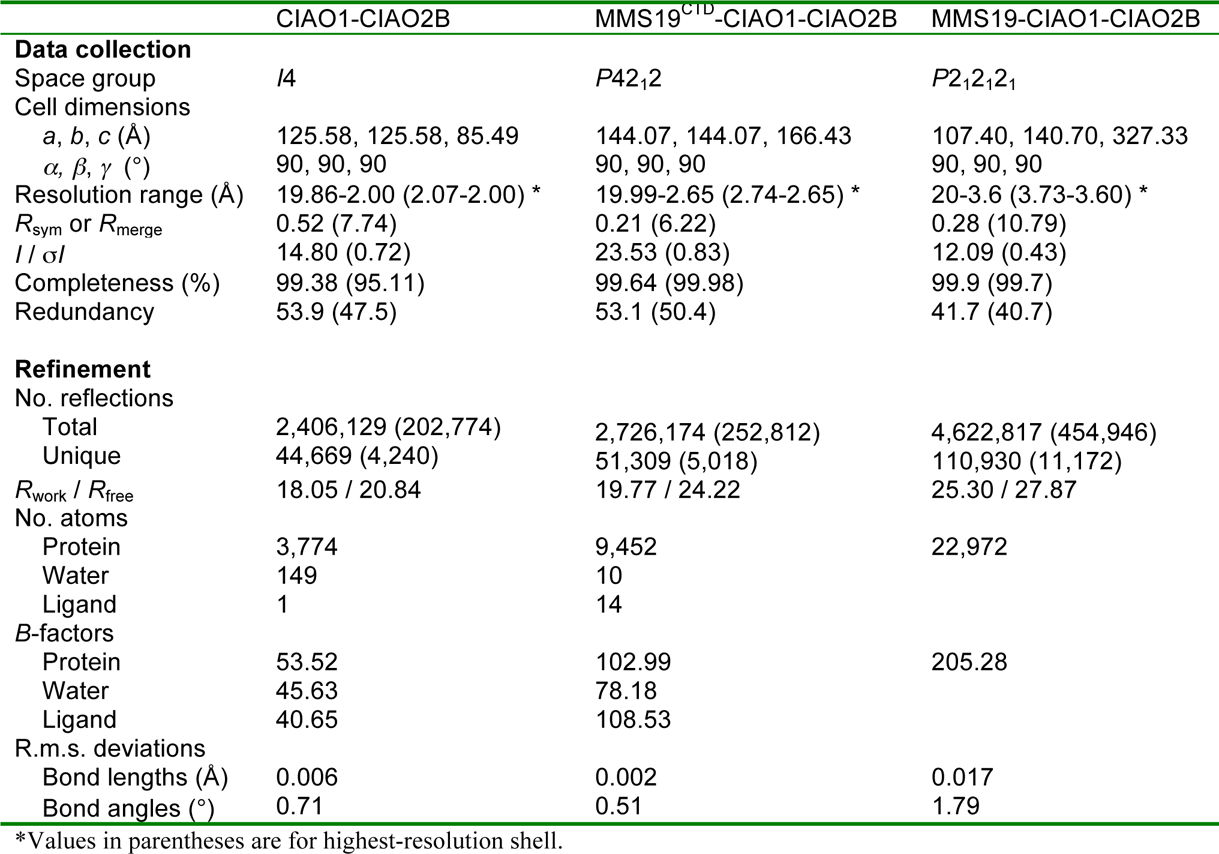
Summary of data collection and refinement statistics of the CIAO1-CIAO2B, MMS19^CTD^-CIAO1-CIAO2B, and MMS19-CIAO1-CIAO2B crystal structures.

To assess the functional conservation of the CIAO1–CIAO2B interaction across species, we used a *Saccharomyces cerevisiae* strain in which expression of the endogenous *CIA1* gene is repressed in the presence of glucose. The resulting growth defect can be rescued by complementation with a plasmid containing a human CIAO1 transgene^10^. Complementation with the same set of mutants employed in our pull-down assay revealed that most residues lining the central cavity of the CIAO1 β-propeller narrow-end face are essential for yeast viability (Extended Data Fig. 1f). An additional conserved patch is found on the side of CIAO1 blade 3 (Extended Data Fig. 1b), in close proximity to Cys86 of CIAO2B (human Cys93^CIAO2B^). This residue was previously identified in a proteomic screen for highly reactive cysteines, is strictly required for yeast survival^24^, and has been implicated in direct Fe-S cluster coordination and transfer^29^. Together, these observations suggest that the conserved patch on blade 3 of CIAO1 might mediate the recruitment of Fe-S client proteins and/or the Fe-S donor CIAO3^19, 30^.

### MMS19 interaction with the core complex

Association of the CIAO1-CIAO2B core complex with the adaptor protein MMS19, which is required for the maturation of DNA replication and repair proteins, was shown to require the four C-terminal HEAT-like repeats in MMS19^21^. To visualize the interaction of MMS19 with the core complex, we crystallized a complex consisting of a C-terminal fragment of mouse MMS19 (residues 911-1031, hereafter referred to as MMS19^CTD^) and *Drosophila* CIAO1-CIAO2B, and determined its structure at a resolution of 2.7 Å (Table 1). The interaction between the catalytic CIAO1-CIAO2B module and the MMS19 adaptor is mediated by the most C-terminal MMS19 HEAT-like repeat (helices α45 and α46), CIAO2B helices α1 and α2, and the loop connecting α2 and β3 (Fig. 1d, e). There are no direct contacts between MMS19 and CIAO1, and CIAO2B acts as a bridge. A pull-down experiment using lysates from HEK293 cells expressing Flag-tagged wild-type and mutant human MMS19 and myc-tagged human CIAO1 and CIAO2B (Fig. 1f and Extended Data Fig. 2b) confirmed that the interaction requires helix α46 of MMS19. Two positively-charged patches in MMS19 were also critical for the interaction: K1007/K1008/K1009^MMS19^, in the loop connecting helices α45 and α46, and R1012/K1013^MMS19^ within helix α46 (Fig. 1f and Extended Data Fig. 2b). These residues were tested in a yeast complementation assay. While yeast MMS19 knock-out strains are viable, they are sensitive to hydroxyurea (HU), a DNA-replication inhibitor^10^. Sensitivity was not restored by expression of MMS19 deletion constructs that lacked helix α46, both helices α45 and α46, or by the charge-reversal mutant R1013E/K1014E^MMS19^ (human R1012/K1013^MMS19^). The effect was less pronounced for the mutant K1008E/K1009E/R1010E^MMS19^ (human K1007/K1008/R1009^MMS19^, Extended Data Fig. 2c). Together, these results confirm that the molecular architecture of the CTC is conserved across eukaryotes.

**Fig. 2.**
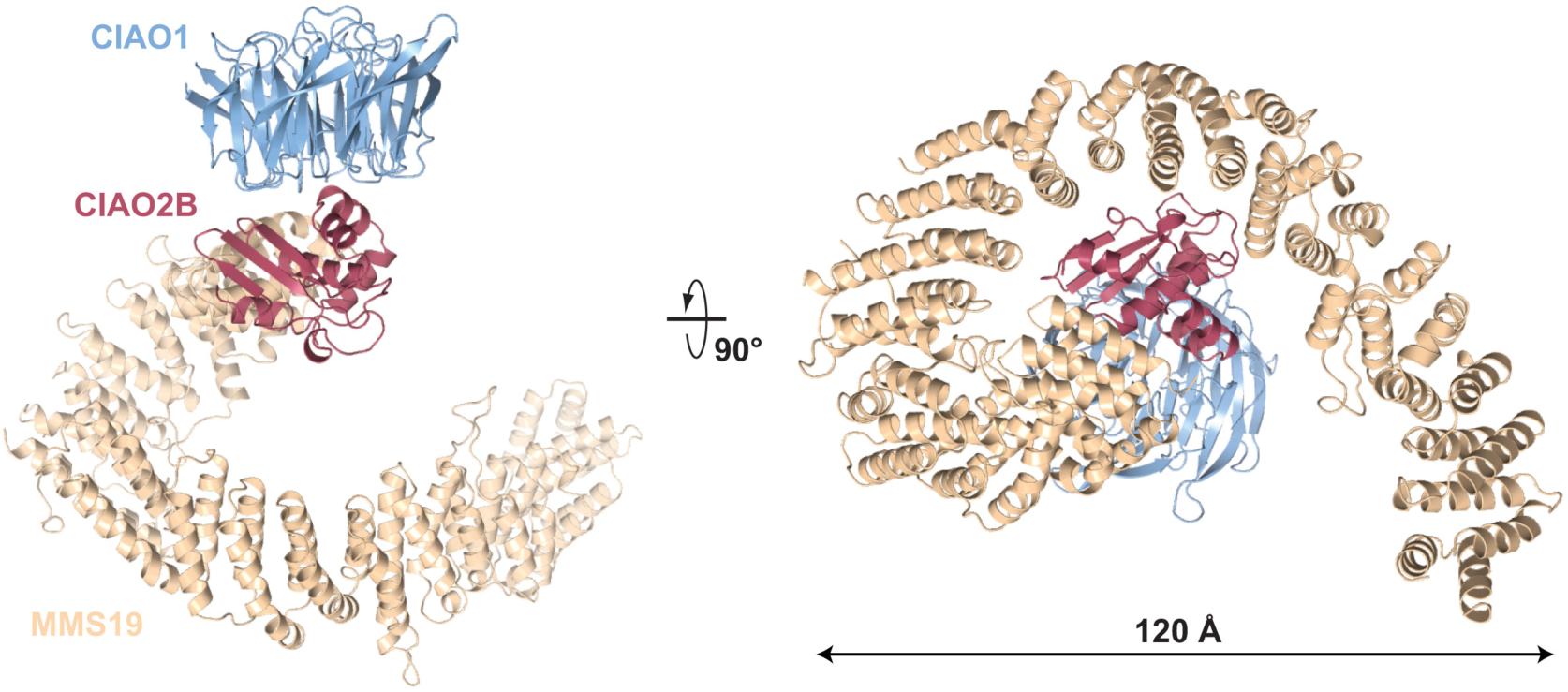
Architecture of the full-length CIA targeting complex. Crystal structure consisting of mouse MMS19 and Drosophila CIAO1-CIAO2B, shown in two orthogonal views.

Helices α45 and α46 of MMS19 were previously shown to be a target for ubiquitination by the MAGE-F1-NSE1 E3 ubiquitin ligase^31^, and ubiquitination was blocked by mutating a cluster of five lysine residues (human K993, K1002, K1007, K1008, K1013) to arginine. Based on our structure, ubiquitination of the MMS19^CTD^ at any of these five positions is likely to preclude CIAO2B binding due to steric clashes (Extended Data Fig. 2d). We therefore expect that ubiquitination of MMS19 is incompatible with binding to CIAO1-CIAO2B, thus preventing formation of the CTC. In addition to targeting MMS19 to the proteasome for degradation, leading to the eventual shut-down of the CIA pathway, modification by ubiquitination might therefore provide a regulatory mechanism that immediately inhibits Fe-S protein biogenesis and prevents further transfer of Fe-S clusters into MMS19-dependent client proteins.

### Architecture of the CIA targeting complex

To understand the architecture of the complete CTC, we determined the crystal structure of a complex comprising full-length mouse MMS19 and *Drosophila* CIAO1-CIAO2B at a resolution of 3.6 Å (Fig. 2). MMS19 consists of 46 α−helices arranged in 23 tandem HEAT-like repeats, which form an overall left-handed super-helical structure, positioning its N-terminus ∼70 Å away from both its C-terminus and the catalytic cysteine of CIAO2B. The structure confirms that CIAO2B interacts with MMS19 solely via the last C-terminal HEAT-like repeat. The remainder of MMS19 is thus accessible for interactions with client proteins or early-acting factors of the CIA pathway.

The MMS19-CIAO1-CIAO2B crystal structure has a dimeric CTC arrangement, mediated by the two MMS19 molecules wrapped around each other with an interface area of 2,950 Å^2^, and an extensive interface between the two CIAO2B protomers (1,644 Å^2^ buried surface area, Extended Data Fig. 2h). Interestingly, CTC dimerization positions potential Fe-S cluster coordinating residues C86, H85, and H119 in close proximity, hinting at how Fe-S cluster coordination might be achieved by the CTC (Extended Data Fig. 2i). CTC dimerization is also observed in solution, as indicated by the co-existence of monomeric and dimeric species in an all-mouse CTC sample analyzed by negative-stain electron microscopy (Extended Data Fig. 2j).

To gain initial insights into how client proteins might be recognized, we mapped evolutionary conservation of amino acid residues on the surface of the CTC, identifying two highly conserved surfaces patches: one in blade 3 of the CIAO1 β-propeller, and the other within the N-terminal domain (NTD) of MMS19 (Extended Data Fig. 2f). Notably, these two patches flank the presumed catalytic cysteine of CIAO2B. These findings suggest that the CTC contains two binding sites for recruitment of Fe-S client proteins: one located on the side of the β-propeller domain of CIAO1, which might represent a general binding site for all Fe-S client proteins, and a second site within the MMS19 NTD involved in recognizing Fe-S client proteins whose biogenesis depends on the MMS19 adaptor (Extended Data Fig. 2g). Alternatively, the CIAO1 site might be required for interactions with the Fe-S carrier CIAO3, while the MMS19 site might be used solely for recruitment of client proteins.

### Cooperative binding of client proteins

To assess the contributions of these conserved sites towards recruitment of client proteins, we employed isothermal titration calorimetry (ITC) to analyze the interaction between CTC and primase, a DNA-dependent RNA polymerase consisting of a catalytic subunit (PriS) and a regulatory subunit (PriL) containing a Fe-S cluster. The equilibrium dissociation constant between full-length CTC and primase was 35 nM (Fig. 3a, Extended Data Table 1). In the absence of MMS19, primase binding affinity for the CIAO1-CIAO2B core complex was reduced to 5.4 μM (Fig. 3b). Without PriL, the PriS subunit still bound to a CIAO1-CIAO2B core complex with an affinity of 3.1 μM (Fig. 3c), indicating that CIAO1 does not directly recognize the primase Fe-S domain. Primase affinity for the MMS19-CIAO2B sub-complex was below the level of detection (Fig. 3d). CTC complexes containing N-terminally deleted MMS19 (constructs MMS19^381–1030^ or MMS19^710–1030^) displayed substantially reduced affinity for primase (508 and 832 nM, respectively, Fig. 3e-f). Collectively, these observations demonstrate that the interaction with the MMS19 NTD increases the affinity of CTC for primase, and indicate that the bipartite binding mode results in a high-affinity, cooperative interaction. This is consistent with previous studies showing that the additional MMS19 binding surface is necessary for the maturation of DNA replication and repair proteins, while the CIAO1 binding site is sufficient for certain clients such as the metabolic enzyme GPAT^10, 28^.

**Fig. 3.**
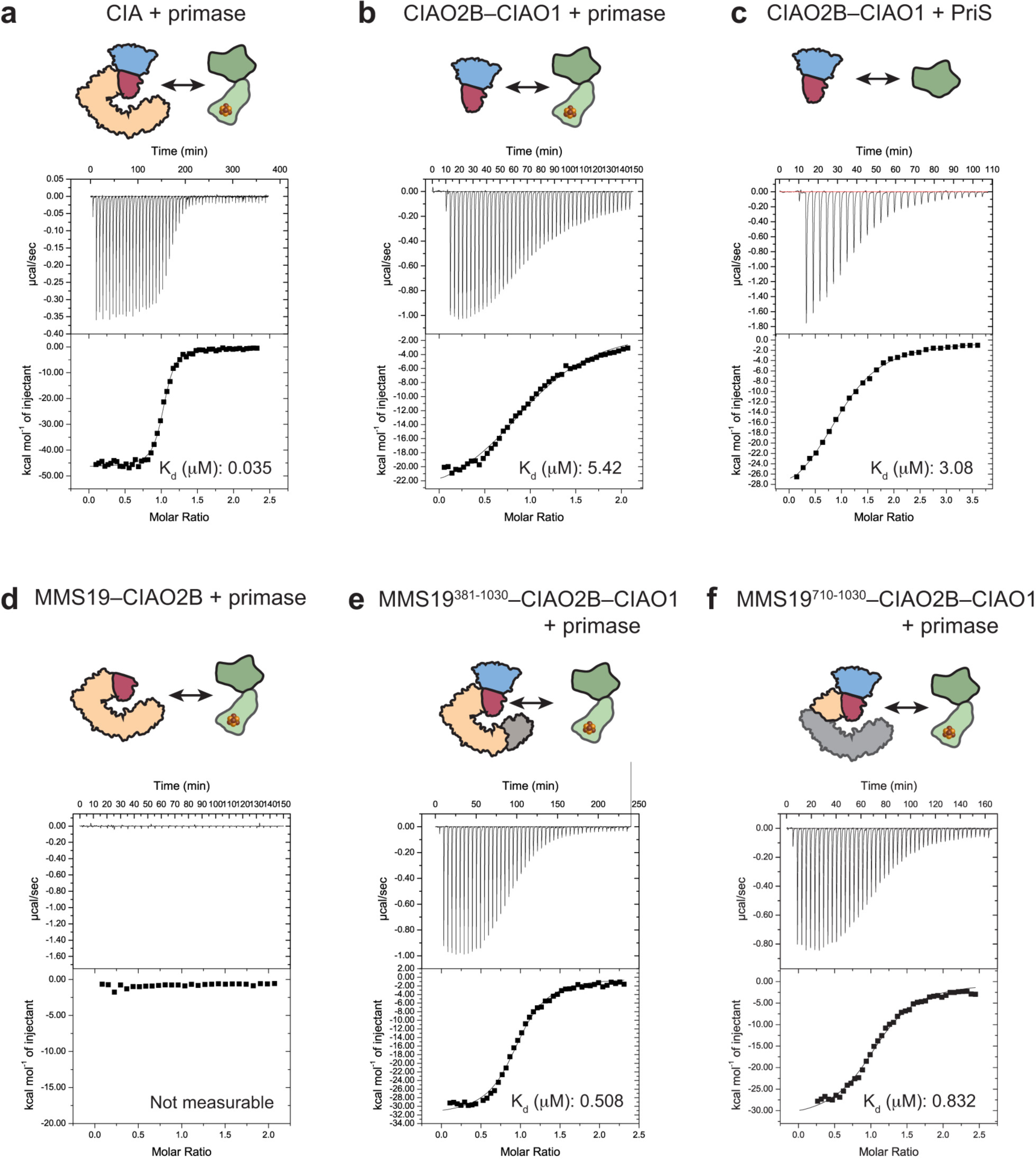
The CIA targeting complex employs cooperative binding via two binding sites on CIAO1 and MMS19 for client protein recognition. ITC binding studies of full-length human CTC and its sub-complexes to full-length client protein primase, or its small subunit PriS. The upper part of each panel shows representative raw ITC data and the lower part shows integrated heat changes. K_d_s estimated from the fitted curves are shown as an inset. Interaction between primase and full-length CIA targeting complex consisting of MMS19–CIAO1–CIAO2B (**a**), the catalytic core module consisting of CIAO1–CIAO2B (**b**), CIAO1–CIAO2B and PriS (**c**), MMS19– CIAO2B and primase (**d**), MMS19^381-1030^–CIAO1–CIAO2B (**e**), and MMS19^710-1030^– CIAO1–CIAO2B (**f**).

### Cryo-EM reconstructions of CTC-client complexes

To directly visualize CTC-client protein interactions, we used single-particle cryo-EM to resolve the architecture of two client protein complexes in which human CTC was bound to the primase PriL-PriS or the DNA helicase DNA2. 2D classification revealed substantial conformational flexibility within MMS19 for both complexes (Fig. 4a-c, Extended Data Fig. 4 and 5). Sorting of particles by 2D and 3D classification followed by multi-body refinement resulted in reconstructions at resolutions of 8.8 Å for CTC-primase, and 11.7 Å for CTC-DNA2 (Fig. 4d-e), which enabled unambiguous docking of the three CTC subunits in both complexes. For the CTC-primase reconstruction, the crystal structure of PriS (PDB 4RR2) was docked into the density directly adjacent to CIAO1, while the N- and C-terminal domains of PriL were docked into the remaining density. For the CTC-DNA2 reconstruction, the precise orientation of DNA2 could not be resolved given its globular shape. Nevertheless, both reconstructions confirm that the two CTC interaction sites in CIAO1 blade 3 and the MMS19 NTD are engaged in client protein binding.

**Fig. 4.**
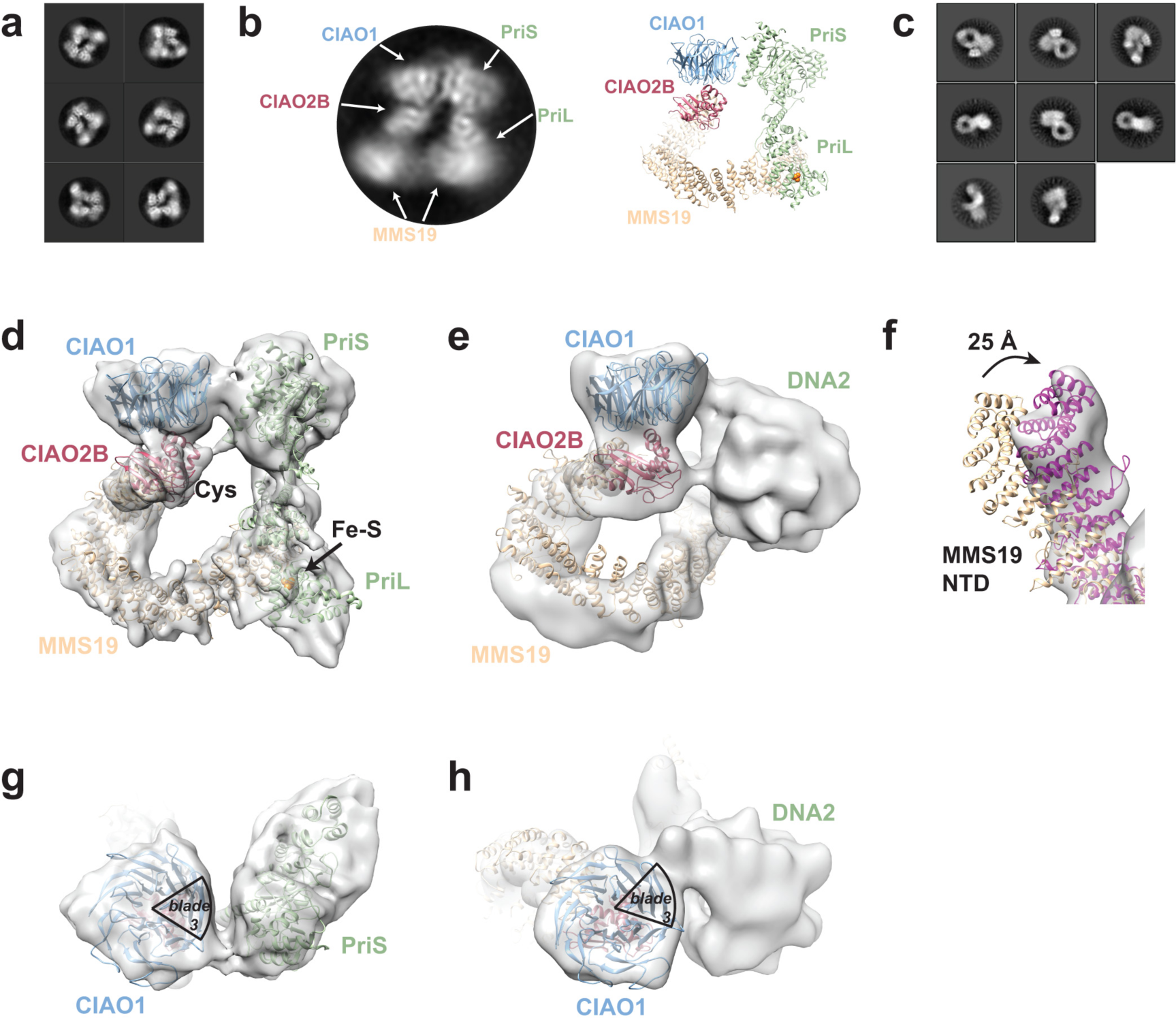
**Conformational plasticity of the CIA targeting complex enables recognition of diverse client proteins. a**, Cryo-EM 2D class averages of CTC– primase. **b**, For comparison, a model of CTC–primase with labelled subunits is shown in cartoon representation. **c**, 2D class averages of CTC–DNA2. **d**, Reconstruction of CTC–primase at 8.8 Å with docked crystal structures of MMS19, CIAO2B, CIAO1, PriS and PriL. **e**, Reconstruction of CTC–DNA2 at 11.7 Å with docked crystal structures of MMS19, CIAO2B, and CIAO1. **f**, View from the bottom of the CTC–DNA2 reconstruction with rigid-body docked MMS19 illustrates the structural rearrangement necessary to accommodate the more globular DNA2 client protein. **g**-**h** View from the top of the CTC–primase (**g**) and CTC–DNA2 (**h**) reconstruction; the location of blade 3 in CIAO1 is indicated.

**Fig. 5.**
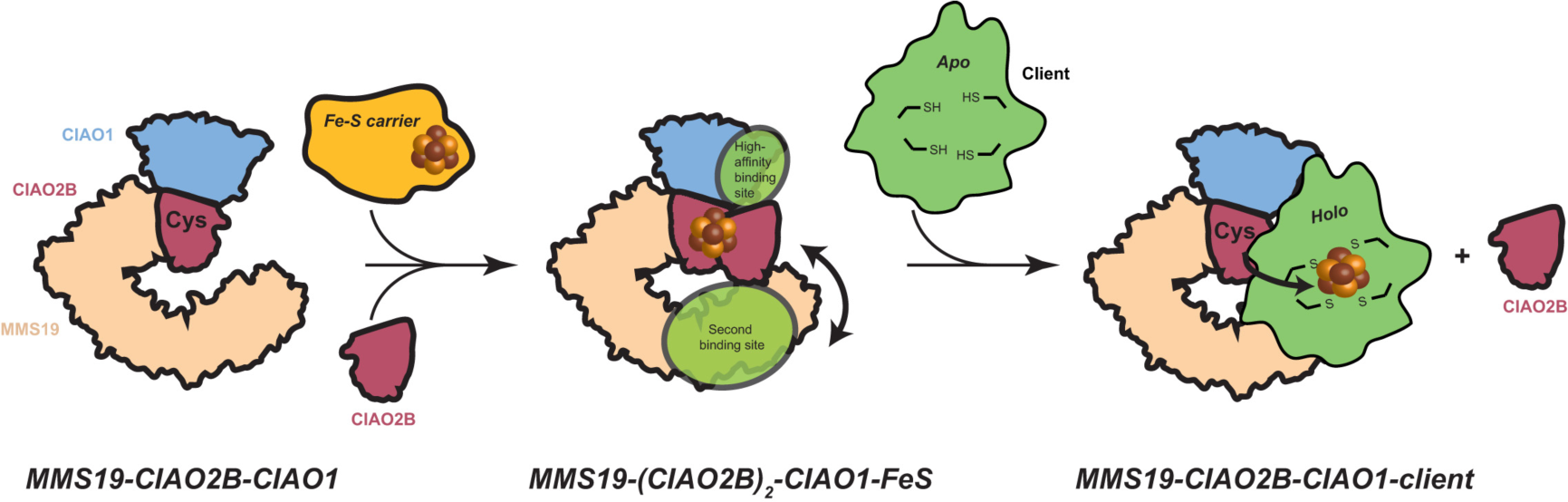
Model of Fe-S cluster transfer and client protein binding by the CTC. The Fe-S cluster is initially transferred from the Fe-S carrier to the CTC, likely involving a second molecule of CIAO2B for stable coordination of the Fe-S cluster (MMS19-CIAO1-(CIAO2B)_2_-FeS). Upon dissociation of the Fe-S carrier, a client protein is recruited via the high-affinity binding site on CIAO1 and the second binding site on MMS19. The Fe-S cluster is then transferred from CIAO2B to the client, potentially aided by the highly reactive cysteine of CIAO2B. After transfer of the Fe-S cluster, the second CIAO2B molecule dissociates, resulting in the MMS19-CIAO1-CIAO2B-client complex observed in the cryo-EM reconstructions.

Analysis of the CTC-primase and CTC-DNA2 reconstructions reveals that both primase and DNA2 interact with CIAO1 blade 3 (Fig. 4g-h). DNA2 contacts CIAO1 within the conserved patch of blade 3 located towards the narrow-end face of the β-propeller (Extended Data Fig. 1), whereas the charged patch located towards the wide-end face of the β-propeller appears unoccupied (Extended Data Fig. 6b). This is consistent with HEK293 cells pull-down assays, which indicate that mutations in this part of CIAO1 do not abolish DNA2 binding (Extended Data Fig. 6a). In contrast, charge reversal mutations in CIAO1 blade 3 prevent PriL-PriS binding to the CTC (Extended Data Fig. 6). Mutation R125E^CIAO1^, which reduces viability in yeast ^26^, abolished the interaction with both primase and DNA2.

Notably, comparisons of the CTC-primase and CTC-DNA2 reconstructions reveal that while the conformation of MMS19 in the CTC-primase complex is similar to its arrangement in the CTC crystal structure, MMS19 adopts a distinct conformation in the CTC-DNA2 structure, with the NTD exhibiting a tighter curvature within its helical repeats (Fig. 4f). This conformational change is necessary to accommodate the more globular DNA2 protein, as compared to the elongated PriL-PriS heterodimer. Our cryo-EM reconstructions therefore suggest that conformational plasticity of MMS19 is important for client protein recognition, and likely responsible for the ability of the CTC to recognize and stably interact with diverse client proteins.

In agreement with our ITC data, the CTC-primase reconstruction confirms that the client protein binding site on CIAO1, which is directly adjacent to the proposed catalytic cysteine of CIAO2B, is occupied solely by the PriS subunit and does not directly recognize the client protein Fe-S domain. The Fe-S cluster-containing PriL subunit, on the other hand, interacts with the MMS19 NTD. In this arrangement, the

Fe-S cluster is positioned ∼70 Å away from Cys93^CIAO2B^, an unexpected finding with implications for the mechanism of Fe-S cluster transfer, as discussed below. Given the observed conformational flexibility of MMS19, the CIA targeting complex might undergo rearrangements to mediate Fe-S cluster transfer from CIAO2B into the client protein, or alternatively accommodate the binding of a Fe-S carrier protein to act as bridge between CIAO2B and the client Fe-S domain.

Altogether, the structures of CTC-client protein complexes demonstrate considerable conformational plasticity of MMS19, which not only facilitates client protein recognition but also likely contributes to the mechanism of Fe-S cluster transfer from CIAO2B into the client.

## Discussion

The targeted insertion of Fe-S clusters into cytosolic and nuclear proteins is a highly regulated process that relies on the CIA pathway and constitutes an essential part of their biogenesis. Despite its critical function in the biogenesis of key DNA replication and DNA repair proteins, the molecular mechanism of the CTC has remained unclear.

Here we used an integrative structural and biochemical approach to uncover the molecular architecture of the central component of the CIA pathway at atomic resolution, and to visualize its interactions with client proteins. Our structural analysis reveals that the CTC contains two interaction sites for recognition of client proteins, a high-affinity binding site on CIAO1, which is likely shared among all client proteins, and a second site within the MMS19^NTD^, which facilitates the recruitment of DNA repair and replication proteins by increasing their binding affinity due to cooperativity between the two sites. Strikingly, the CTC-primase cryo-EM reconstruction reveals that the high-affinity binding site on CIAO1 interacts with the catalytic PriS subunit, which is located ∼70 Å away from the Fe-S cluster in the PriL subunit. Thus, our structural analysis reveals that at least for some client proteins, CIAO1 does not directly recognize their Fe-S domains or any structural motifs proximal to the Fe-S cluster. This binding mode is likely universal, given that prior interaction studies on XPD showed that the CTC binding site resides within its arch domain and not within the Fe-S domain^32^.

Comparisons of our cryo-EM reconstructions of CTC in complexes with DNA2 or PriL-PriS revealed that the conformational flexibility of MMS19 plays an important role in accommodating diverse clients. While the CIAO1-CIAO2B catalytic core represents a rigid unit, the plasticity of the alpha-helical subunit MMS19 allows the CTC to adapt to diverse molecular shapes of client proteins, and potentially other co-factors required for Fe-S cluster transfer. Thereby, MMS19 bears a striking similarity to other α-helical repeat proteins that rely on structural flexibility to mediate protein-protein interactions in cellular processes ranging from protein (de)phosphorylation to ubiquitination and nuclear transport^33–36^. Karyopherins, HEAT-repeat proteins that recognize diverse macromolecular cargoes and mediate their transport between the nucleus and the cytoplasm, convert between two functionally and conformationally distinct nuclear and cytoplasmic states, which facilitate cargo binding and release^34, 37^. The HEAT-repeat comprising A subunit of protein phosphatase 2A, a Ser/Thr phosphatase involved in cell cycle regulation, undergoes an extensive conformational rearrangement that moves its NTD by ∼60 Å upon interaction with the regulatory B subunit^33^. Similar parallels can be drawn between the CTC and other molecular machineries involved in protein modification, such as cullin-like E3 ubiquitin ligases, which undergo conformational changes to enable recognition of protein substrates and to facilitate ubiquitin transfer^35^. In the context of the CIA pathway, the structural plasticity of the CTC is likely not only critical for its ability to recognize diverse client proteins, but may also facilitate large-scale (>50 Å) molecular motions necessary for Fe-S cluster transfer and incorporation.

Our findings provide critical structural insights that have implications for the Fe-S cluster transfer mechanisms of the CTC. The MMS19-CIAO1-CIAO2B crystal structure has a dimeric CTC arrangement with an extensive interface between CIAO2B protomers (Extended Data Fig. 2h), and analysis by negative stain EM indicates that the CTC exists in a monomer-dimer equilibrium in solution (Extended Data Fig. 2j). In the homologous CIAO1-CIAO2A system, which is specifically required for the maturation of aconitase, a CIAO1-(CIAO2A)_2_ heterotrimer was shown to coordinate a Fe-S cluster under anaerobic conditions^29^. A similar mode of cluster coordination is likely to occur in the CIAO1-CIAO2B system, as supported by the oligomerization states observed in our structural studies. We propose a mechanism whereby the Fe-S cluster is transferred from an Fe-S carrier protein onto a MMS19-CIAO1-(CIAO2B)_2_ complex (Fig. 5). Although the dimeric holo-CTC is incompatible with client protein recruitment due to steric clashes between the client and the second MMS19 molecule, an additional CIAO2B molecule can be readily accommodated by client-bound CTC (Extended Data Fig. 7). Given the distal arrangement of the client Fe-S domain and the catalytic cysteine involved in cluster transfer, CIAO2B dimerization would shield the Fe-S cluster until a client protein is bound. Transfer can then occur either directly across the CTC, relying on MMS19 conformational flexibility, or possibly by involving a Fe-S carrier protein such as CIAO3, which might additionally contribute to cluster stability. Upon Fe-S cluster transfer, CIAO2B would then be released, resulting in the MMS19-CIAO1-CIAO2B-client product complex observed in our cryo-EM reconstructions. Our model is in agreement with proteomic studies that showed an excess of CIAO2B over CIAO1 and MMS19 in HeLa cells (176 nM MMS19, 455 nM CIAO1, 1253 nM CIAO2B)^38^ and in mouse fibroblasts^39^. The proposed transfer mechanism thus likely exploits the compositional and conformational dynamics of the CTC to facilitate client apo-protein recognition, Fe-S cluster loading, insertion, and finally Fe-S protein release.

In conclusion, our structural and functional studies of the CTC visualize the central component of the CIA machinery and reveal a critical role of conformational dynamics of its MMS19 subunit for client protein recognition and likely also Fe-S cluster transfer.

## METHODS

No statistical methods were used to predetermine sample size. The experiments were not randomized. The investigators were not blinded to allocation during experiments and outcome assessment.

### Protein expression and purification

Full-length human, mouse and Drosophila MMS19, CIAO2B and CIAO1 were sub-cloned into pAC-derived vectors^40^, adding either a Strep II or a hexa-histidine tag to their N-terminus, followed by a tobacco etch virus (TEV) protease cleavage site.

Point mutants were generated using the Quikchange Site-directed Mutagenesis Kit (Stratagene) according to the manufacturer’s protocol. Baculoviruses were produced through recombination in *Spodoptera frugiperda* (Sf9) insect cells (unauthenticated and untested for mycoplasma contamination, Thermo Fisher Scientific B82501) grown in EX-CELL 420 medium (Sigma) by co-transfection of the pAC-derived vector containing the gene of interest and viral DNA^41^ using Cellfectin II Reagent (Thermo Fisher Scientific). Following virus amplification in Sf9 insect cells, recombinant CIA complexes and subunits were expressed in *Trichoplusia ni* High-Five insect cells (unauthenticated and untested for mycoplasma contamination, Expression Systems 94-002F) infected at a density of 2.0 x 10^6^ ml^-1^ with the appropriate virus or combination of viruses, and grown in Sf-900 II medium (Thermo Fisher Scientific) at 27 **°**C and 120 rpm. Cells were harvested after 40-48 hours, resuspended in buffer containing 50 mM Tris pH 8.0, 200 mM NaCl, 1 mM TCEP, 2 mM PMSF, and Complete EDTA-free protease inhibitor tablets (Roche), and lysed by sonication. The lysate was clarified by ultracentrifugation at 40,000 g for 40 min. The supernatant was then filtered through Miracloth (Merck) and passed once through an empty Econo-Pac chromatography column (Bio-Rad) before being loaded onto a column packed with the appropriate affinity chromatography matrix.

#### MmMMS19-DmCIAO1-CIAO2B

Full-length mouse MMS19, and *Drosophila melanogaster* CIAO2B and CIAO1 were sub-cloned into pAC-derived vectors^40^, adding a Strep II-tag to the N-terminus of MMS19 and a hexa-histidine tag to the N-termini of CIAO2B and CIAO1. The protein complexes were purified using Ni-NTA agarose resin (Qiagen), resulting in a mixture of MMS19-CIAO2B-CIAO1 and CIAO2B-CIAO1 complexes. His- and Strep-tags were removed by cleavage with TEV protease during overnight dialysis against buffer containing 50 mM Tris pH 8.0, 100 mM NaCl, 1 mM TCEP. The complex was further purified by Poros HQ ion exchange chromatography (Thermo Fisher Scientific) and a subtractive Ni-NTA affinity step. Trimeric MMS19-CIAO2B-CIAO1 was separated from dimeric CIAO1-CIAO2B by size exclusion chromatography on a Superdex 200 column (GE Life Sciences) in 20 mM HEPES pH 7.4, 200 mM NaCl, 1 mM TCEP. The two peaks were pooled separately and concentrated to 27 mg/ml (MMS19-CIAO2B-CIAO1 complex) and 35 mg/ml (CIAO1-CIAO2B complex).

#### HsMMS19-CIAO2B-CIAO1 and its subcomplexes and truncations

Human MMS19-CIAO2B-CIAO1, its subcomplexes, truncations, and individual subunits for ITC measurements and electron microscopy analysis, were expressed recombinantly in High-Five insect cells using the baculovirus expression system, and purified as described above for the *Mm*MMS19-*Dm*CIAO2B-CIAO1 complex by Ni-NTA affinity, Poros HQ ion exchange chromatography, and size exclusion chromatography. Prior to ITC measurements, samples were purified on a Superdex 200 column (GE Life Sciences) in assay buffer (20 mM HEPES pH 7.4, 150 mM NaCl, 3% glycerol, 0.5 mM TCEP).

#### MmDNA2

Full-length mouse DNA2 was sub-cloned into a pAC-derived vector, adding a hexa-histidine tag followed by a TEV cleavage site to its N-terminus. The protein was recombinantly expressed in High-Five insect cells using the baculovirus expression system and purified by Ni-NTA affinity chromatography. DNA2-containing fractions were pooled and further purified on a heparin column (HiTrap, GE Life Sciences), followed by size exclusion chromatography on a Superdex 200 column (GE Life Sciences) in buffer containing 20 mM HEPES pH 7.4, 100 mM NaCl, and 1 mM TCEP.

#### HsPriL-PriS and HsPriS

Human primase was expressed and purified as described^42, 43^, with minor modifications. For expression of PriL-PriS, plasmids pBG100-p48 and pETDuet-p58 (a kind gift from W. Chazin) were transformed into BL21 Star (DE3) cells (Thermo Fisher). The culture was grown in TB medium at 37**°**C and the temperature was lowered to 18 **°**C when the cell density approached OD_600_ of 0.6. Expression was induced with 1 mM IPTG and continued for 20 hours at 18 **°**C. Upon induction, the culture medium was supplemented with 0.1 mg/ml ferric citrate. Cells were harvested, resuspended in buffer containing 50 mM Tris pH 8.0, 150 mM NaCl, 3% glycerol, 1 mM TCEP, 2 mM PMSF, Complete EDTA-free protease inhibitor tablets (Roche) and snap frozen in liquid nitrogen. Cells were lysed using an Emulsiflex C3 homogenizer (Avestin). The lysate was cleared by ultracentrifugation and by passing it through a 5 μm filter before being loaded onto a Ni-NTA affinity column (Qiagen). The eluate was further purified on a heparin column and a Superdex 200 column (GE Life Sciences), eluting in buffer containing 20 mM HEPES pH 7.4, 150 mM NaCl, 3% glycerol, and 1 mM TCEP. For expression of human PriS, plasmid pBG100-p48 was transformed into BL21 Star (DE3) cells (Thermo Fisher). The culture was grown in TB medium as described above for the PriL-PriS complex. Expression was induced with 1 mM IPTG and continued for 20 hours at 18 **°**C. Cells were harvested, resuspended and lysed, and PriS was purified as described above for the PriL-PriS complex.

### Crystallization and data collection

Crystallization experiments were performed in MRC 2-well crystallizations plates (Swissci) with a Phoenix dispensing robot (Art Robbins) using the sitting-drop vapor diffusion method at 20 **°**C.

X-ray diffraction data were collected at beam lines X06DA (PXIII) and X10SA (PXII) of the Swiss Light Source (Paul Scherrer Institute, Villigen, Switzerland) at a wavelength of 1 Å (unless noted otherwise below) from crystals cooled to 100 K. Diffraction data were processed using XDS or autoproc^44, 45^. Data collection and refinement statistics are summarized in Extended Data Table 1.

#### DmCIAO1-CIAO2B

Crystals of Drosophila CIAO1-CIAO2B were grown in 400 nl hanging drops containing equal volumes protein solution at 35 mg/ml and a reservoir solution consisting of 0.1 M Tris pH 8.5 and 25% PEG MME 2k. Crystals appeared within hours, grew to their final size within 2 days, and were harvested after 7 days and flash cooled in liquid nitrogen.

#### MmMMS19^CTD^-DmCIAO1-CIAO2B

Crystals of a chimeric complex composed of mouse MMS19^CTD^ (residues 911-1031) and the *Drosophila* CIAO1-CIAO2B complex were grown in 600 nl hanging drops containing 400 nl protein solution at 7.8 mg/ml and 200 nl reservoir solution consisting of 0.2 M KSCN and 20% PEG 3350. Crystals appeared after 4 days and were harvested after 6 days using 20% ethylene glycol as a cryo-protectant.

#### MmMMS19-DmCIAO1-CIAO2B

Crystals of a chimeric complex composed of mouse MMS19 and the *Drosophila* CIAO1-CIAO2B complex were grown in 600 nl hanging drops containing 400 nl protein solution at 27 mg/ml and 200 nl reservoir solution consisting of 100 mM HEPES pH 7.5 and 25% PEG 3350. Crystals appeared after 2 days, grew to their final size within 4 days, and were harvested after 6 days. Crystals were cryo-protected in a buffer containing 100 mM HEPES pH 7.5, 30% PEG 3350, and 10% ethylene glycol, and flash cooled in liquid nitrogen.

#### MmMMS19-MmCIAO1-CIAO2B

For experimental phasing, crystals of the all-mouse *Mm*MMS19-*Mm*CIAO1-CIAO2B complex were grown in 600 nl hanging drops containing 400 nl protein solution at 12 mg/ml and 200 nl reservoir solution consisting of 100 mM HEPES pH 7.0, 207 mM ammonium citrate, 21% PEG 3350, and 0.5% jeffamine ED-2001. For a native sulfur single wavelength anomalous dispersion (S-SAD) experiment, data were collected at 5975 eV (2.0751 Å). For phasing, crystals were derivatized by soaking in mother liquor supplemented with hexatantalum tetradecabromide (Jena Bioscience) prior to harvesting. Diffraction data were collected at the Ta L-III peak absorption edge (9880.5 eV, 1.255 Å, f’ = −17.27, f’’ = 23.25).

### Crystal structure determination and refinement

Crystals of the *Dm*CIAO1-CIAO2B complex diffracted to 2.0 Å and belonged to the tetragonal space group I4. The structure was solved by molecular replacement in Phaser, as implemented in Phenix^46, 47^, using the structure of human CIAO1 as a search model (PDB 3FM0,^27^). Model building was performed in Coot^48^. The final model was refined in Phenix to an *R*_work_ factor of 18.05% and an *R*_free_ factor of 20.84%, and contains residues 2-17 and 32-155 of CIAO2B, and 2-335 of CIAO1 (Table 1). The asymmetric unit contained one copy of the *Dm*CIAO1-CIAO2B heterodimer, with residues 2-17 of CIAO2B packing against a crystallographic symmetry-related copy of CIAO2B. Analysis with Molprobity^49, 50^ shows 98.5% of all residues within favoured regions of the Ramachandran plot with one outlier.

Crystals of the *Mm*MMS19^CTD^-*Dm*CIAO1-CIAO2B complex diffracted to 2.7 Å and belonged to the tetragonal space group P42_1_2. The structure was determined by molecular replacement in Phaser as implemented in Phenix^46, 47^, using the structure of the Drosophila CIAO1-CIAO2B complex as a search model. The asymmetric unit contains two copies of the *Mm*MMS19^CTD^-*Dm*CIAO1-CIAO2B complex. Iterative model building and refinement were carried out with the programs Coot and Phenix. The model was refined to an *R*_work_ factor of 19.77% and an *R*_free_ factor of 24.22% (Table 1), and contains residues 2-21 and 28-155 of CIAO2B, residues 2-335 of CIAO1, and residues 913-1031 of MMS19. Analysis with Molprobity shows 97.9% of all residues within favoured regions of the Ramachandran plot with no outliers.

Crystals of the *Mm*MMS19-*Dm*CIAO1-CIAO2B and the *Mm*MMS19-*Mm*CIAO1-CIAO2B complex belonged to the orthorhombic space group P2_1_2_1_2_1_ and contained two copies of the MMS19-CIAO2B-CIAO1 complex in the asymmetric unit. Best diffraction was obtained from a crystal of *Mm*MMS19-*Dm*CIAO1-CIAO2B complex to 3.6 Å.

The structure was solved by single-wavelength anomalous dispersion (SAD) phasing using a dataset obtained from Ta_6_Br_12_-derivatized crystals of the *Mm*MMS19-*Mm*CIAO1-CIAO2B complex. Heavy atom search was performed in SHELXD and phasing was carried out in SHARP^51, 52^. The resulting phases were improved by density modification with Parrot as implemented in ccp4^53^. The resulting map revealed clear density corresponding to MMS19, into which ideal poly-serine α-helices were placed. The model was then refined against using deformable elastic network (DEN) restraints in CNS solve^54, 55^, which led to major improvement of the maps, enabling placement of CIAO2B, CIAO1 and the *Mm*MMS19^CTD^ fragment using the higher-resolution *Dm*CIAO1-CIAO2B and *Mm*MMS19^CTD^-*Dm*CIAO1-CIAO2B structures.

The resulting model was completed by iterative building and refinement against the native *Mm*MMS19-*Dm*CIAO1-CIAO2B crystal data. The amino acid sequence of MMS19 was assigned based on bulky density visible for large hydrophobic side chains, as well as information on the location of cysteine and methionine residues determined using native sulfur SAD phasing data obtained from a crystal of the *Mm*MMS19-*Mm*CIAO1-CIAO2B complex. Further iterative cycles of refinement and model building were carried out initially with autoBUSTER^56^, applying NCS restraints, target restraints to the MMS19^CTD^-CIAO1-CIAO2B structure applied as LSSR, and TLS refinement. Subsequent refinement in Phenix, using NCS and target restraints as well as TLS refinement, produced a final model with an *R*_work_ factor of 25.3% and an *R*_free_ factor of 27.9% (Extended Data Table 1). The final model contains residues 2-155 of CIAO2B, 2-335 of CIAO1, and 15-43, 46-539, 548-638, 649-699 (chain C), 649-696 (chain F), and 702-1028 of MMS19. Analysis with Molprobity shows 95.3% of all residues within favoured regions of the Ramachandran plot with 0.07% outliers.

### Electron microscopy sample preparation and data collection

For negative stain analysis, human MMS19-CIAO2B-CIAO1 complex was mixed with a 1.4-molar excess of mouse DNA2 or human PriL-PriS, incubated on ice for 30 min, and purified on a Superose 6 10/300 column (GE Healthcare) in 20 mM HEPES, 75 mM NaCl, and 1 mM TCEP. The chromatogram and analysis by SDS-PAGE indicated that the complexes were stoichiometric and homogeneous (Extended Data Fig. 4a-b and 5a-b). However, initial characterization of a uranyl-formate stained sample from the peak fraction revealed that the complex was extremely fragile and prone to disintegration on the EM grid. The complex was stabilized by cross-linking on ice for 10 min with 0.5% (final) glutaraldehyde (Ted Pella). Continuous carbon grids (Ted Pella, 01840-F) were glow-discharged for 90 sec at 15 mA using an easiGlow glow discharge cleaning system (PELCO). The sample was diluted to ∼70 nM and 4 μl were incubated on an EM grid for 1 min. After blotting with filter paper, the grids were subsequently laid onto 5 drops of 1% (w/v) uranyl formate solution. Data were collected on a Tecnai Spirit transmission electron microscope (Thermo Fisher) operating at 120 kV with a nominal magnification of 49,000x using EPU (Thermo Fisher), and recorded on a FEI Eagle 4k CCD camera, with a final pixel size of 1.52 Å.

For cryo-EM analysis, the sample was prepared and cross-linked as above, and diluted to 0.3 mg/ml (CTC-primase) or 0.4 mg/ml (CTC-DNA2). 4 μl sample were applied onto an UltraAuFoil gold grid (R 1.2/1.3, Quantifoil) prepared with a thin home-made carbon film floated on top, and glow-discharged using the easiGlow glow discharge cleaning system for 90 s at 15 mA. The sample was incubated on the grid for 30 s, blotted for 3 s at force setting −1, and plunge frozen in liquid ethane using a Vitrobot Mark IV (Thermo Fisher).

Data were collected on a Cs-corrected (CEOS GmbH) Titan Krios transmission electron microscope (Thermo Fisher) operating at 300 kV using EPU (Thermo Fisher). The CTC-primase dataset was recorded at a nominal magnification of 130k on a Falcon II direct electron detector, resulting in a final pixel size of 0.87 Å. Exposures were fractionated into 50 frames, with a total dose of 50 e^-^ Å^-2^. A total of 3,128 movies were acquired at a defocus range from −0.6 to −3 μm. The CTC-DNA2 dataset was recorded using a Volta phase plate on a Gatan K2 summit direct electron detector with a Quantum LS energy filter (slit width 20 eV), at a nominal magnification of 105k, resulting in a final pixel size of 1.1 Å. Exposures were fractionated into 40 frames, with a total dose of 39.9 e^-^ Å^-2^. A total of 1,948 movies were acquired at a defocus range from −0.4 to −0.8 μm.

### Electron microscopy data processing

For negative stain data, particles were picked automatically using DoG picker^57^. Particles were extracted, and 2D classification, initial model generation and 3D refinement performed in SPARX and Relion^58–60^. For cryo-EM data, particles were picked with Gautomatch (http://www.mrc-lmb.cam.ac.uk/kzhang/), using templates based on low-pass filtered 2D class averages from a previous data set collected using a Volta phase plate. Concomitantly with data collection, movies were motion corrected using MotionCor2^61^, and CTF parameters were estimated using Gctf^62^.

Using cryoflare, an in-house software wrapper for on-the-fly processing, data quality was assessed and suitable micrographs were selected for further processing (www.cryoflare.org). 2D and 3D classification, as well as 3D refinement and multi-body refinement were performed in Relion. For multi-body refinement, partially overlapping masks were employed that contained the CIAO1-CIAO2B-DNA2 subcomplex and MMS19-CIAO2B, or the CIAO1-CIAO2B-PriS-PriL and MMS19-CIAO2B subcomplexes, respectively. For details of the data processing scheme, see Extended Data Figure 4 (CTC-primase) and Extended Data Figure 5 (CTC-DNA2). Both CTC-primase and CTC-DNA2 samples showed preferred orientation (Extended Data Fig. 4g and 5g), which could not be overcome in the presence of a range of different detergents, or by glow-discharging in the presence of amylamine.

### ITC measurements

Protein samples were purified on a Superdex 200 column (GE Life Sciences) in assay buffer (20 mM HEPES pH 7.4, 150 mM NaCl, 3% glycerol, 0.5 mM TCEP) prior to ITC measurements. ITC measurements were performed at 25 **°**C using a Microcal VP-ITC Calorimeter (Malvern) with a stirring speed of 307 rpm and a reference power of 10 μCal s^-1^. Experimental parameters for calorimetric titrations of the CIA complex and its subcomplexes (in the syringe) and primase (in the cell) were optimized for each sample. Data were analyzed with MicroCal ITC Origin 7 analysis software (Origin-based software provided by Malvern).

### Pull-down assays in HEK293 cells

HEK Expi 293 cells (unauthenticated and untested for mycoplasma contamination, Thermo Fisher Scientific 14527) were grown in FreeStyle 293 expression medium (Thermo Fisher Scientific) at 37 **°**C, 85% humidity, and 135 rpm. All plasmids used for transfection were based on the GATEWAY system (Invitrogen), contained a cytomegalovirus (CMV) promoter, and expressed either Flag- or myc-tagged protein constructs (see Extended Data Table 3 for details). Cells were seeded at 0.6 x 10^6^ ml^-1^ 24 h prior to transfection. For transfection of 10 ml cells in suspension, 10 μg of DNA were diluted into 400 μl of Opti-MEM reduced serum media (Thermo Fisher Scientific). In a separate tube, 25 μl of polyethyleneimine (Polysciences) at 1 mg/ml were diluted into 400 μl of Opti-MEM reduced serum media and then added to the DNA solution. The transfection mix was added drop-wise to the cells. Cells were harvested after 42 hours, resuspended in a buffer containing 50 mM Tris pH 8.0, 200 mM NaCl, 5% glycerol, 1 mM TCEP, 0.1% Triton-X-100, 2 mM PMSF, Complete EDTA-free protease inhibitor tablets (Roche), and lysed by sonication. Clarified lysates were incubated with Flag-M2 beads (Sigma-Aldrich) for 2 hours at 4 **°**C. After extensive washing, bound proteins were eluted in buffer containing 0.5 mg/ml 1x Flag peptide (Sigma-Aldrich) for 2 hours at 4 **°**C. Samples were analyzed by SDS-PAGE and Western blotting using Anti-Flag M2-peroxidase (HRP) antibody (Sigma-Aldrich), or c-Myc tag antibody (Genescript) and Anti-mouse IgG-peroxidase antibody (Sigma-Aldrich), using Clarity ECL Western Blotting Substrate (Bio-Rad).

### Yeast drop assays

Yeast were grown in YPAD or minimal medium with appropriate selection markers at 30 **°**C. For details of haploid *S. cerevisiae* strains see Extended Data Table 4. Plasmids were transformed into yeast using a standard protocol using lithium acetate, single-stranded carrier DNA and PEG 4000. For drop assays, overnight cultures of yeast cells were diluted to an OD_600_ of 0.7. Five 10-fold serial dilutions were prepared, and 2 μl each were dropped onto selective plates. Plates were incubated at 30 **°**C and imaged after 2-3 days.

### Illustrations and figures

Sequence alignments were generated using Clustal Omega^63^ and colored with ESPript^64^. Structure figures were generated with PyMOL (www.pymol.org) and Chimera^65^.

### Data availability

The atomic coordinates and structure factors reported in this study have been deposited in the Protein Data Bank (PDB) under accession numbers 6TBN (CIAO1-CIAO2B complex), 6TBL (MMS19^CTD^-CIAO1-CIAO2B complex), and 6TC0 (MMS19-CIAO1-CIAO2B complex). The cryo-EM density maps have been deposited in the Electron Microscopy Data Bank (EMDB) under accession codes EMD-XXX (CTC-primase) and EMD-XXX (CTC-DNA2).

## Acknowledgments

We are grateful for technical support from FMI core facilities: S. Cavadini, A. Schenk, A. Graff Meyer, and C. Genoud (electron microscopy), H. Gut, J. Keusch, and G. Kempf (X-ray crystallography), and V. Iesmantavicius, D. Hess and J. Seebacher (mass spectrometry). We are grateful to the staff at beamlines PXII and PXIII of the Swiss Light Source (Paul Scherrer Institute, Villigen, Switzerland) for assistance with X-ray data collection. We thank W. Chazin for providing the plasmids for recombinant expression of human primase in bacteria, and purified primase protein for initial studies; A. Potenza for help with protein expression in the early stage of the project; R. Bunker for help and advice with crystallographic data collection and processing; K. Shimada, M. Hauer and I. Deshpande for sharing protocols and advice on yeast experiments; K. Gari and D. Odermatt for sharing plasmids and helpful discussions; M. Jinek for advice on crystallographic data collection, processing, and model building, and critical reading of the manuscript; J. Reinert and F. Bleichert for comments on the manuscript.

S.K. was supported by a long-term postdoctoral fellowship from the European Molecular Biology Organization (EMBO, ALTF 871-2014). This work was funded by the Swiss National Science Foundation through Sinergia grant number CRSII3_160734, and by the European Research Council under the European Union’s Horizon 2020 Research and Innovation program, grant number 666068, to N.T.

## Author contributions

Conceptualization, S.K. and N.T; Data curation, S.K.; Formal analysis, S.K.; Funding acquisition, S.K. and N.T; Investigation, S.K.; Methodology, S.K.; Project administration, S.K. and N.T; Resources, S.K.; Supervision, N.T.; Validation, S.K. and N.T; Visualization, S.K.; Writing - original draft, S.K.; Writing - review and editing, S.K. and N.T.

## Corresponding author

Correspondence and requests for materials should be addressed to N.T.

## Competing interests

The authors declare no competing interests.

**Supplementary Information** is available for this paper.

## SUPPLEMENTARY INFORMATION

**Extended Data Fig. 1.**
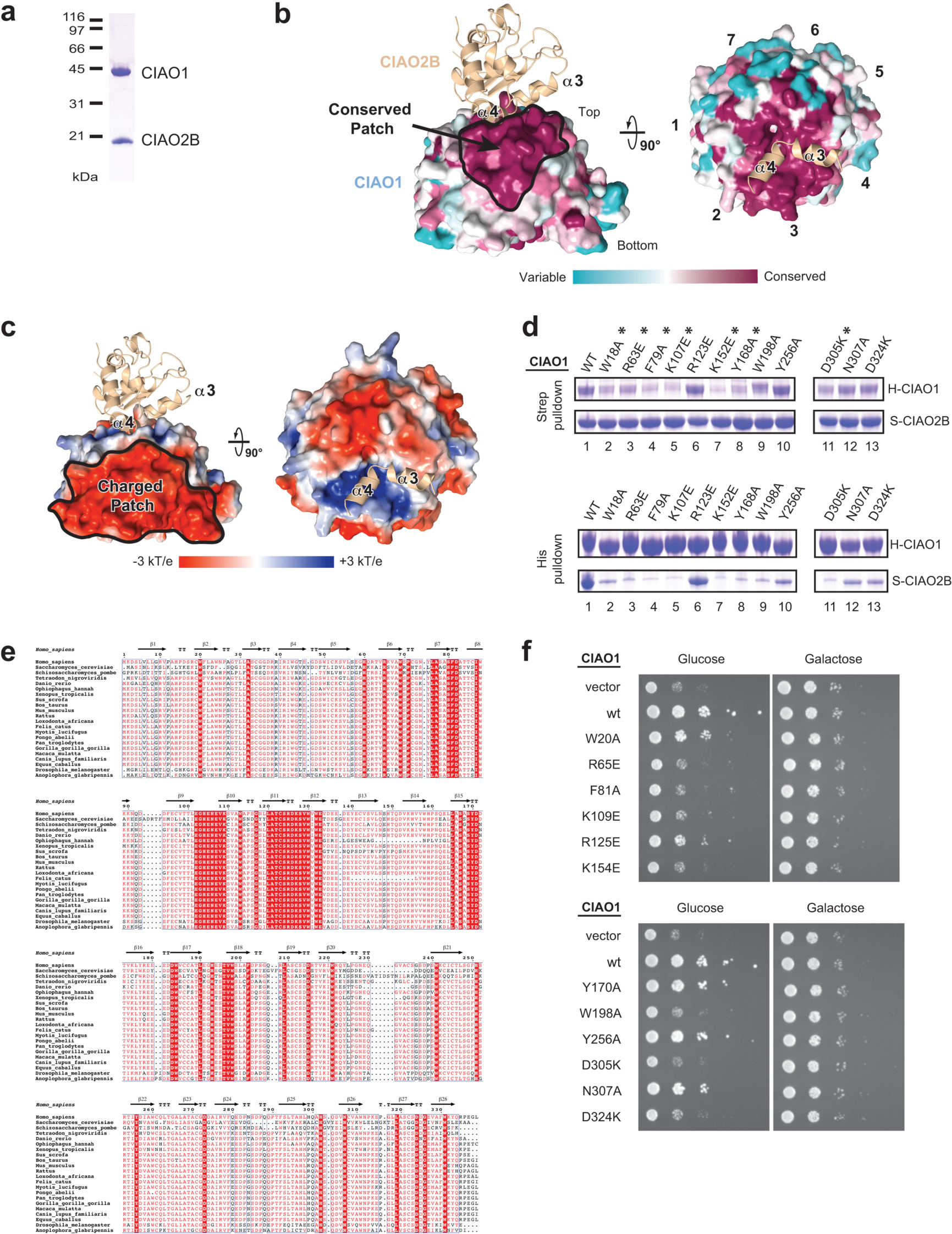
**Crystal structure of the CIAO1-CIAO2B core complex. a**, Coomassie-stained SDS-PAGE of purified *Drosophila melanogaster* CIAO1-CIAO2B complex. **b**, Surface representation of CIAO1 colored according to amino acid conservation across species based on the alignment in **d**. **c**, Surface representation of CIAO1 colored according to electrostatic potential from −3 to +3 k_B_T/e. **d**, CIAO1 sequence alignment used to map evolutionary conservation in **a**. Secondary structure elements of human CIAO1 are indicated above the alignment. Numbering is relative to the human sequence. **e**, Strep pull-down assay from Hi5 insect cells expressing Strep-CIAO2B and His-CIAO1 wt or indicated mutants. **f**, Yeast drop assay of CIAO1 point mutations located at the top face of the beta-propeller. A Gal-CIA1 strain was transformed with either empty vector, wt human CIAO1, or a CIAO1 point mutant; a dilution series of each culture was spotted on agar plates in the presence of glucose or galactose.

**Extended Data Fig. 2.**
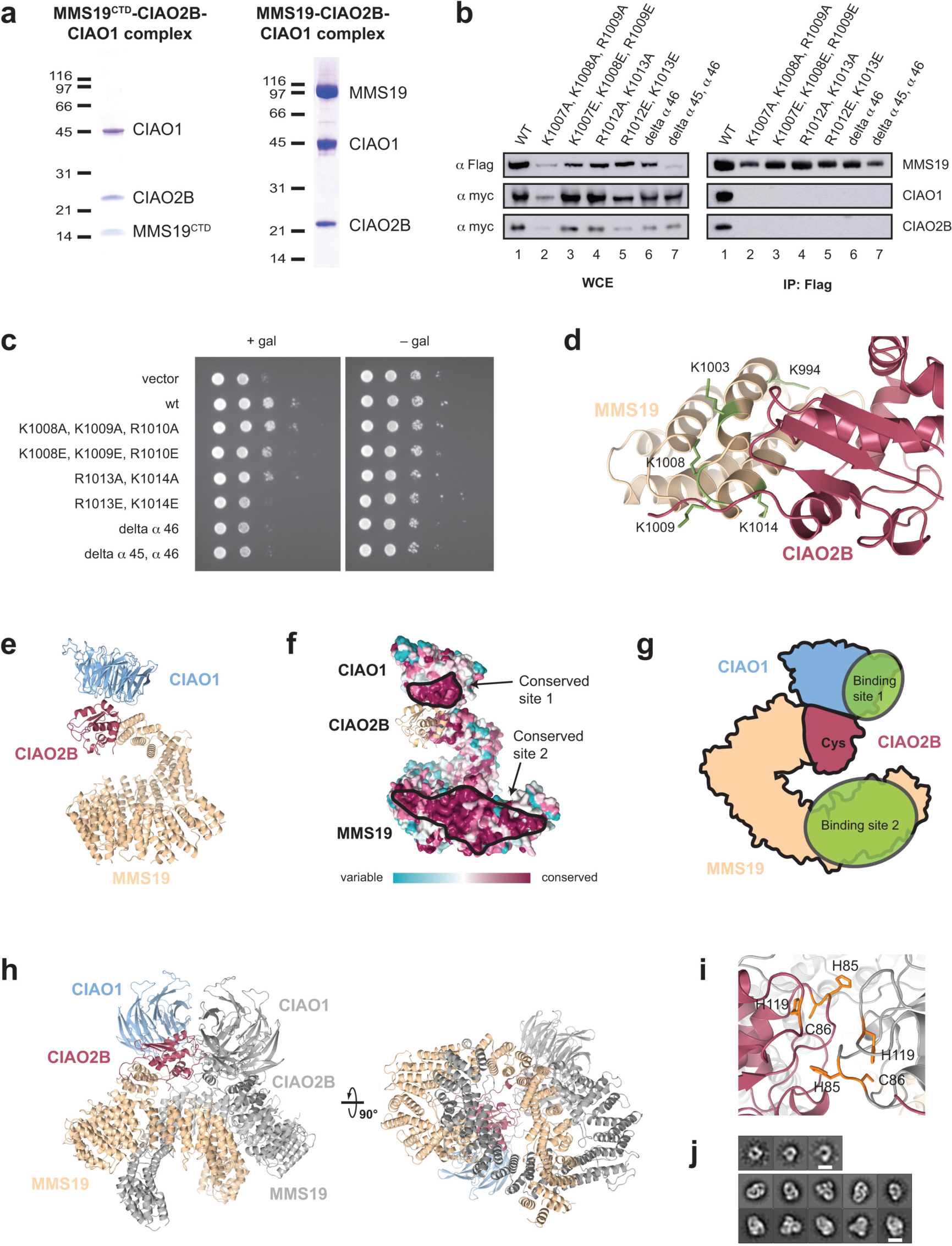
**Crystal structures of the MMS19-CIAO2B-CIAO1 CIA targeting complex. a**, Coomassie-stained SDS-PAGE gels of purified MMS19^CTD^-CIAO2B-CIAO1 and MMS19-CIAO2B-CIAO1 complexes. **b**, Pull-down assay from HEK293 cells using Flag-tagged human MMS19 wt and mutants, and myc-tagged human CIAO2B/CIAO1. WCE (right) and eluted proteins (IP:Flag, left) were analyzed by Western blot. Corresponding to Fig. 1f. **c**, Yeast drop assay of MMS19 mutations. A MMS19 knockout strain was transformed with either empty vector, wt MMS19, or a MMS19 mutant; a dilution series of each culture was spotted on agar plates in the presence or absence of galactose and 20 mM HU. **d**, Lysine residues implicated as targets for ubiquitination by MAGE-F1-NSE1 are shown in green. **e**, Crystal structure of the CIA targeting complex shown in the same orientation as in **f**. **f**, Conservation of amino acid residues across species mapped onto the surface of the CTC structure. **g**, Schematic indicating the location of the two client-protein binding sites as deduced from analysis of the crystal structure of the CIA targeting complex. **h**, Cartoon representation of the dimeric crystal structure of *Mm*MMS19-*Dm*CIAO2B-CIAO1 complex as observed in the crystal. **i**, Close-up view of the CIAO2B-CIAO2B dimer interface. Potential Fe-S cluster-coordinating residues H85, C86, and H119 are shown as sticks. **j**, Reference-free 2D class averages of a negatively stained mouse MMS19-CIAO2B-CIAO1 sample showing monomeric (top) and dimeric (bottom) CIA targeting complexes. *Scale bar represents 100 Å*.

**Extended Data Fig. 3.**
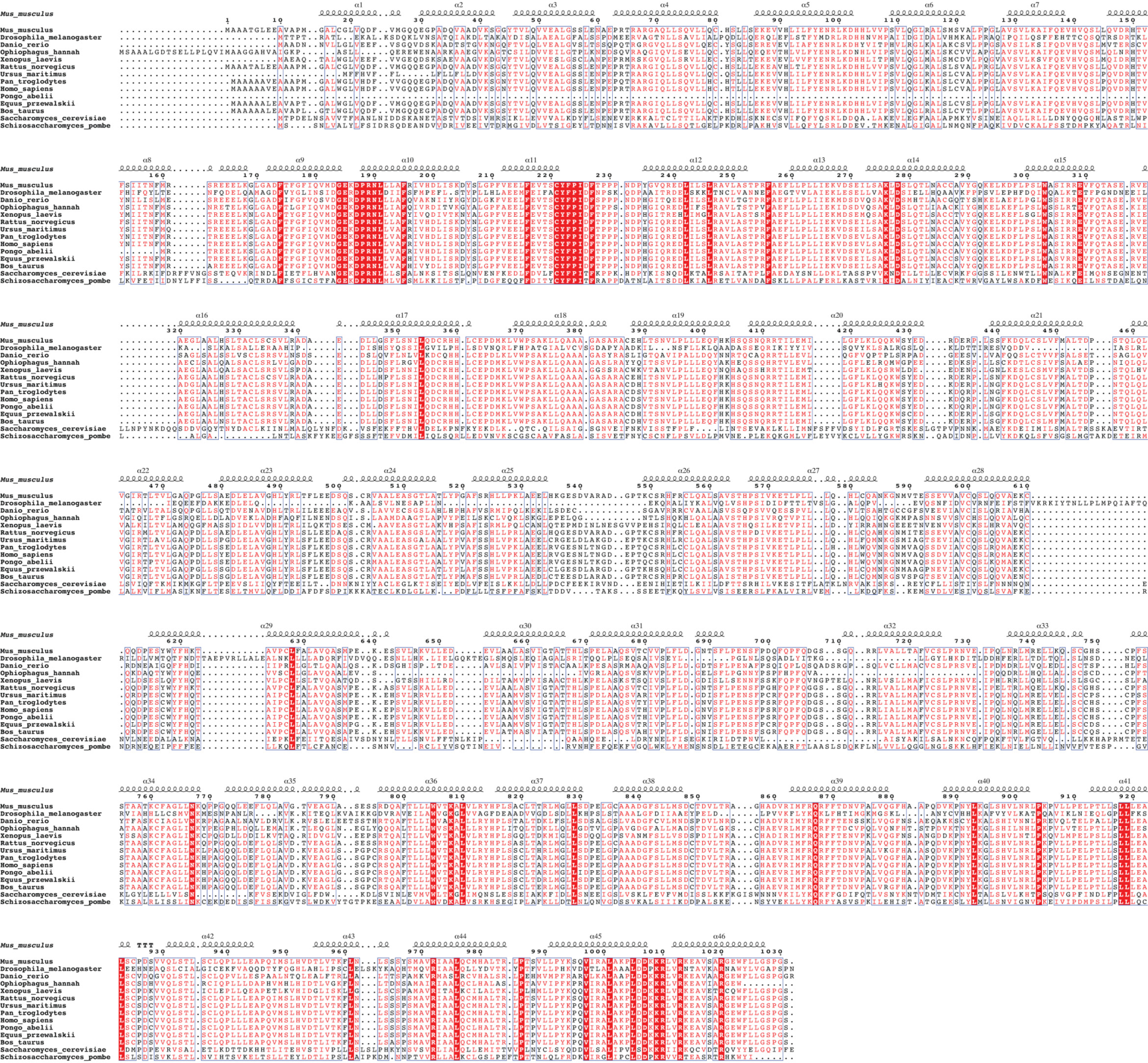
MMS19 sequence alignment used to map evolutionary conservation in Extended Data Fig. 2.

**Extended Data Fig. 4.**
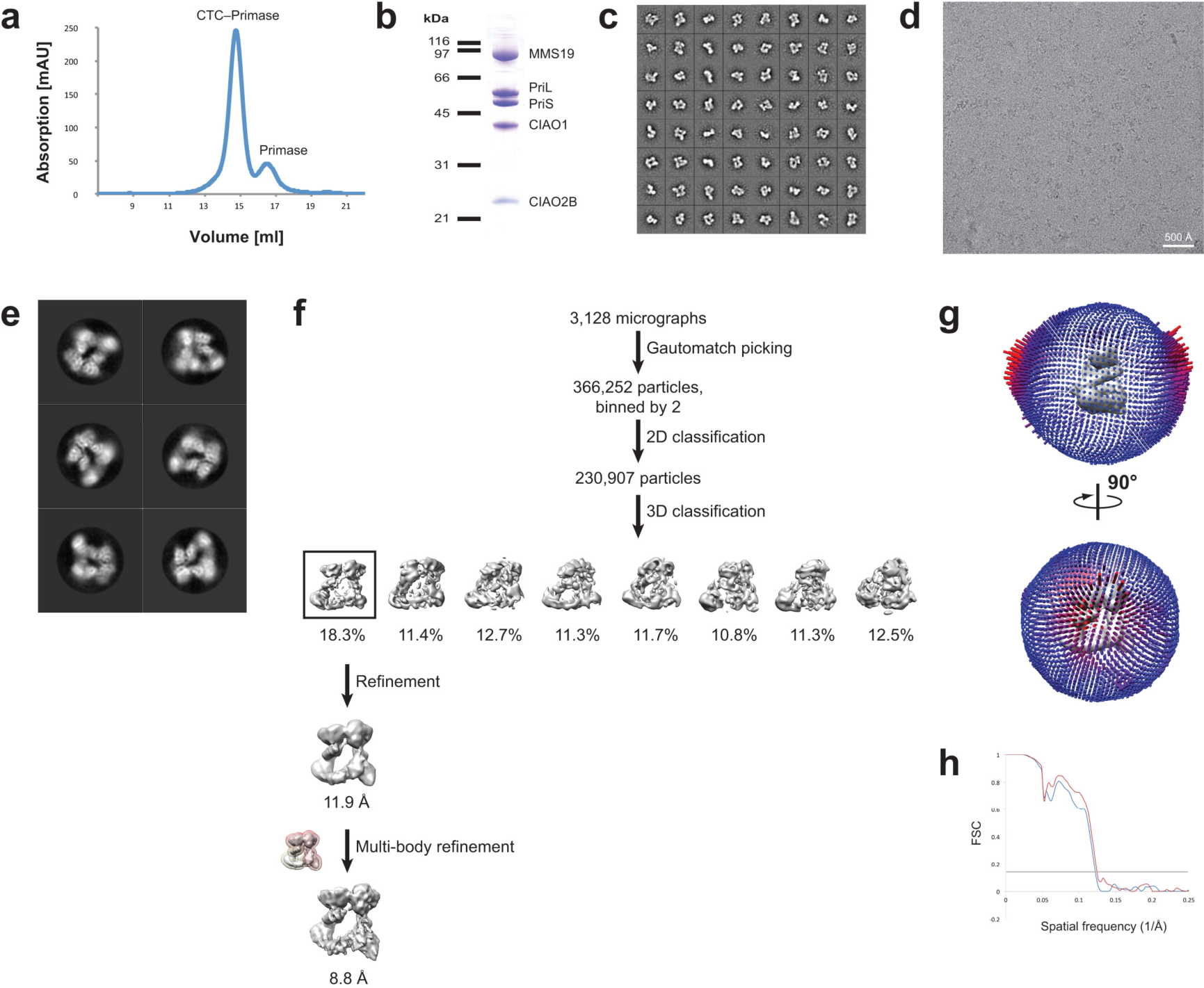
**Cryo-EM reconstruction of a CTC-primase complex. a**, Size exclusion chromatography profile of assembled CTC-primase complex. **b**, Coomassie-stained SDS-PAGE of CTC-primase complex. **c**, Negative stain 2D class averages of the CTC-primase complex. **d**, Cryo-EM micrograph of CTC-primase sample. **e**, Representative 2D class averages of the CTC-primase data set. **f**, Overview of data processing and classification scheme. **g**, Angular distribution of particle orientations in the reconstruction. **h**, FSC plot for half-maps of the reconstruction, 0.143 FSC criterion is indicated.

**Extended Data Fig. 5.**
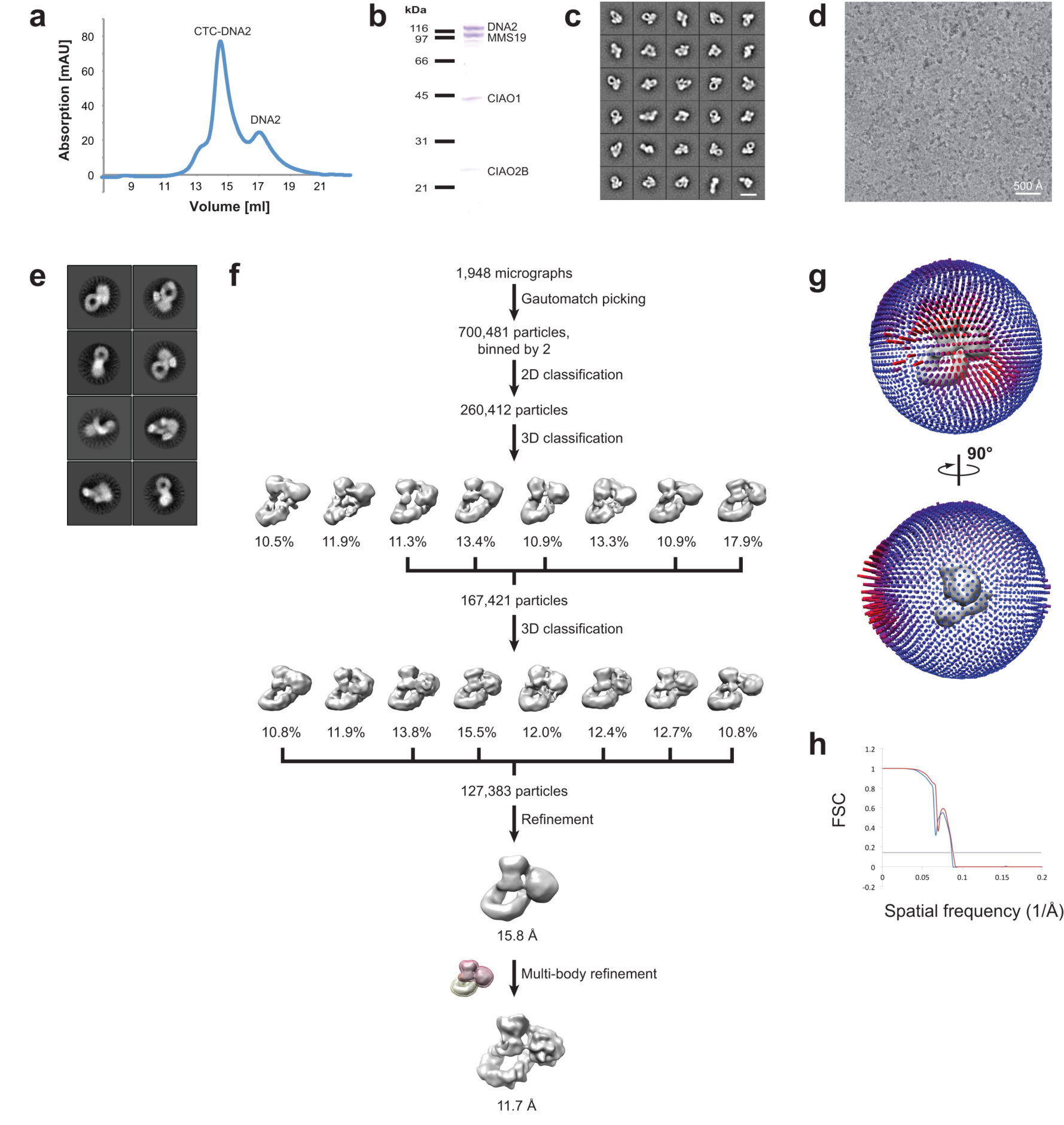
**Cryo-EM reconstruction of a CTC-DNA2 complex. a**, Size exclusion chromatography profile of assembled CTC-DNA2 complex. **b**, Coomassie-stained SDS-PAGE of CTC-DNA2 complex. **c**, Negative stain 2D class averages of the CTC-DNA2 complex. **d**, Cryo-EM micrograph of CTC-DNA2 sample. **e**, Representative 2D class averages of the CTC-DNA2 data set. **f**, Overview of data processing and classification scheme. **g**, Angular distribution of particle orientations in the reconstruction. **h**, FSC plot for half-maps of the reconstruction, 0.143 FSC criterion is indicated.

**Extended Data Fig. 6.**
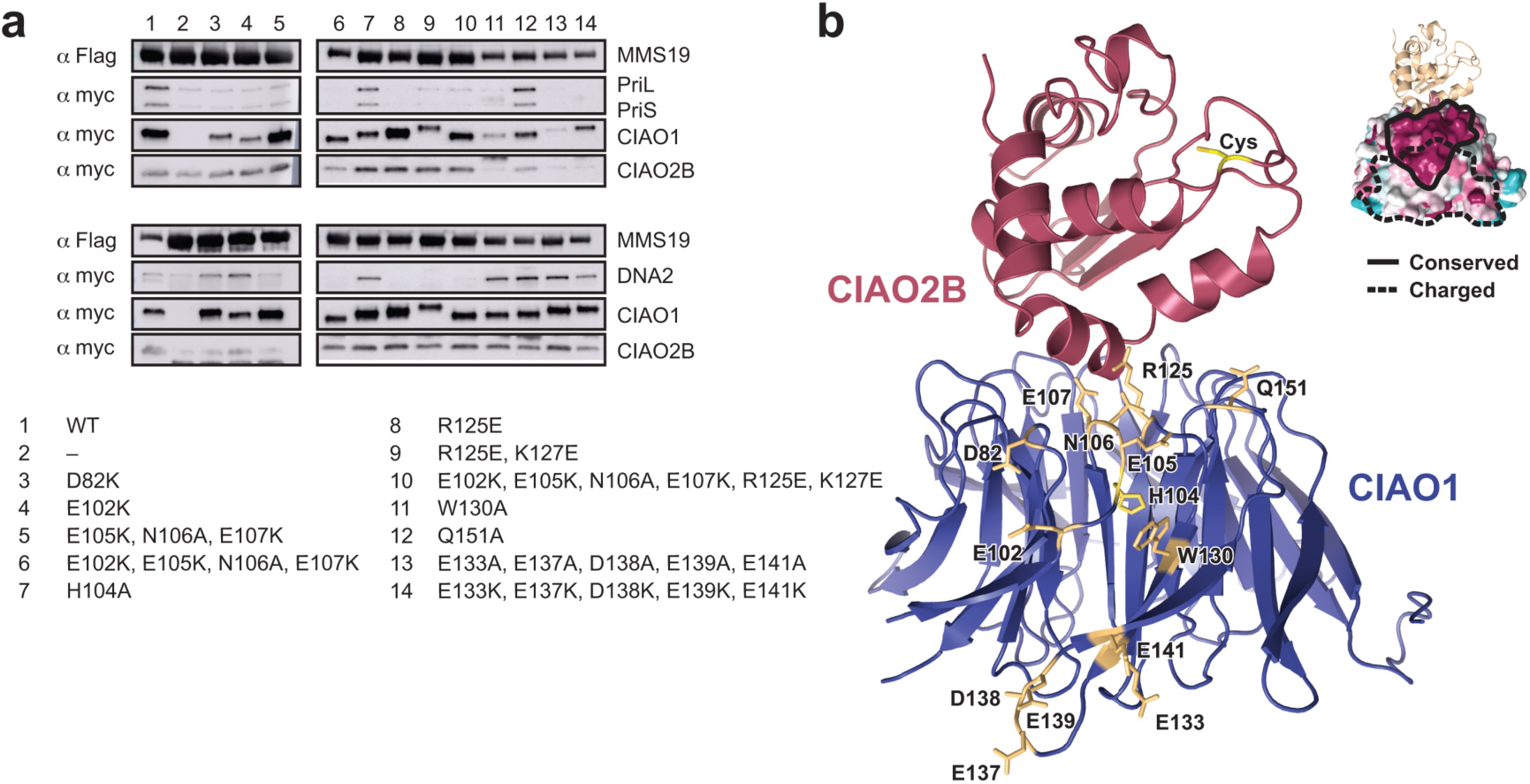
Pull-down assays in mammalian cells pinpoint blade 3 in CIAO1 as the main interaction site for recruitment of client proteins. a, Co-expression and co-IP of Flag-MMS19 with myc-CIAO2B, wild-type or mutant myc-CIAO1, and either myc-PriL/S or myc-DNA2 in HEK293 cells. b, Location of mutated amino acid residues around blade 3 of CIAO1. The inset shows the location of the conserved and charged patches in blade 3 of CIAO1.

**Extended Data Fig. 7.**
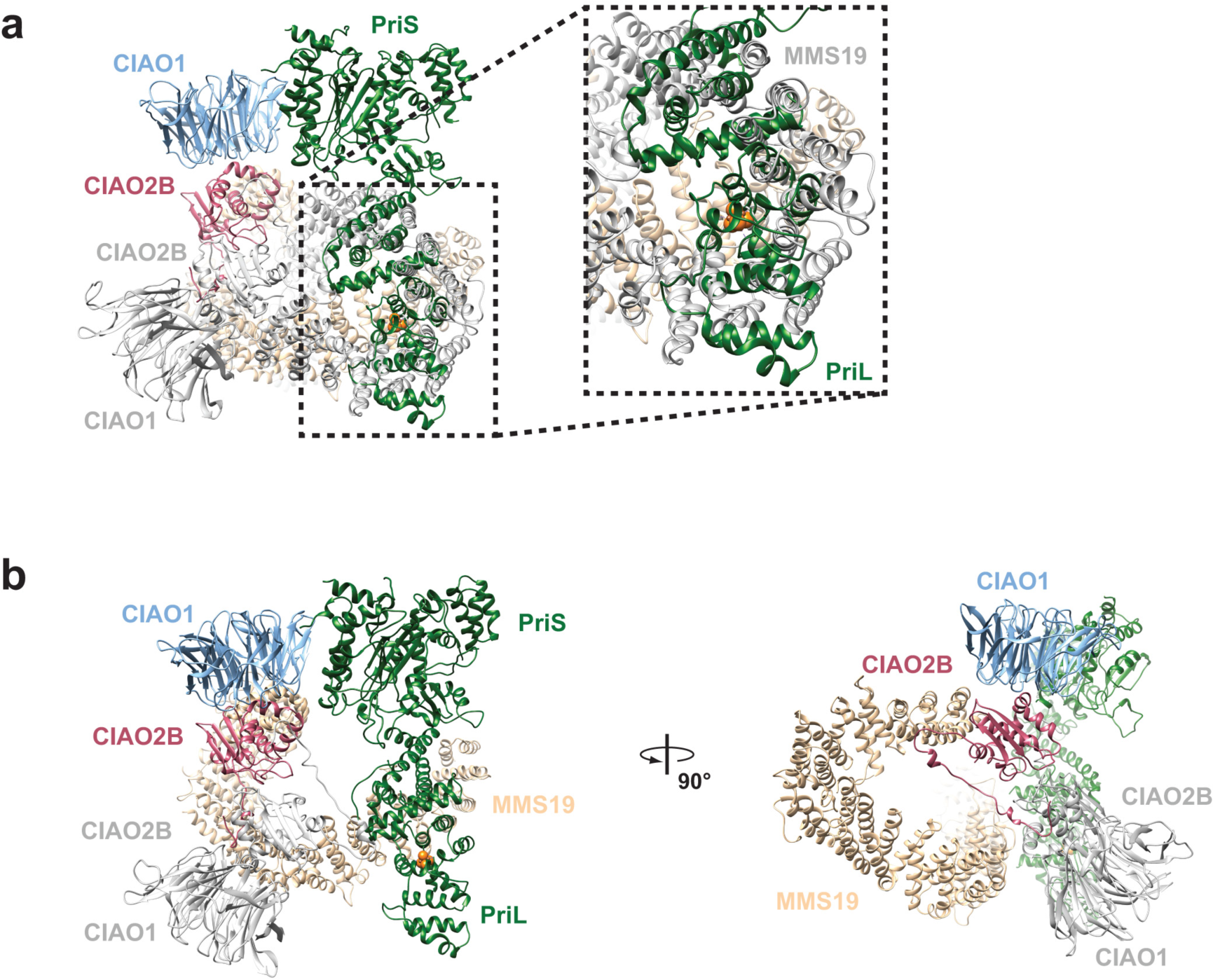
**Superposition of the CTC dimer found in the crystal structure and the CTC-primase complex observed by cryo-EM. a**, Superposition of the CTC dimer with the CTC-primase model results in significant steric clashes between the second MMS19 molecule (grey) and the large subunit of primase. **b**, Superposition of the CIAO1-CIAO2B complex shows that the catalytic core can be readily accommodated in between the CTC and primase without any steric clashes.

**Extended Data Table 1.** Summary of isothermal titration calorimetry experiments.

| Syringe | Cell | K<sub>d</sub><br>[μM] | N | Δ H<br>[kcal/mol] | Δ S<br>[cal/mol/deg] |
| --- | --- | --- | --- | --- | --- |
| MMS19-CIAO1-CIAO2B | PriS-PriL | 0.04 | 1.02 | - 46.5 | - 122 |
| CIAO1-CIAO2B | PriS-PriL | 4.3 | 1.08 | - 25.4 | - 60.9 |
| CIAO1-CIAO2B | PriS | 3 | 0.95 | - 33.23 | - 86.2 |
| MMS19-CIAO2B | PriS-PriL | n.d. | n.d. | n.d. | n.d. |
| MMS19<sup>381-1030</sup>-CIAO1-CIAO2B | PriS-PriL | 0.56 | 0.97 | - 32.16 | - 79.3 |
| MMS19<sup>710-1030</sup>-CIAO1-CIAO2B | PriS-PriL | 1.08 | 1.11 | - 32.25 | - 80.9 |

**Extended Data Table 2.**
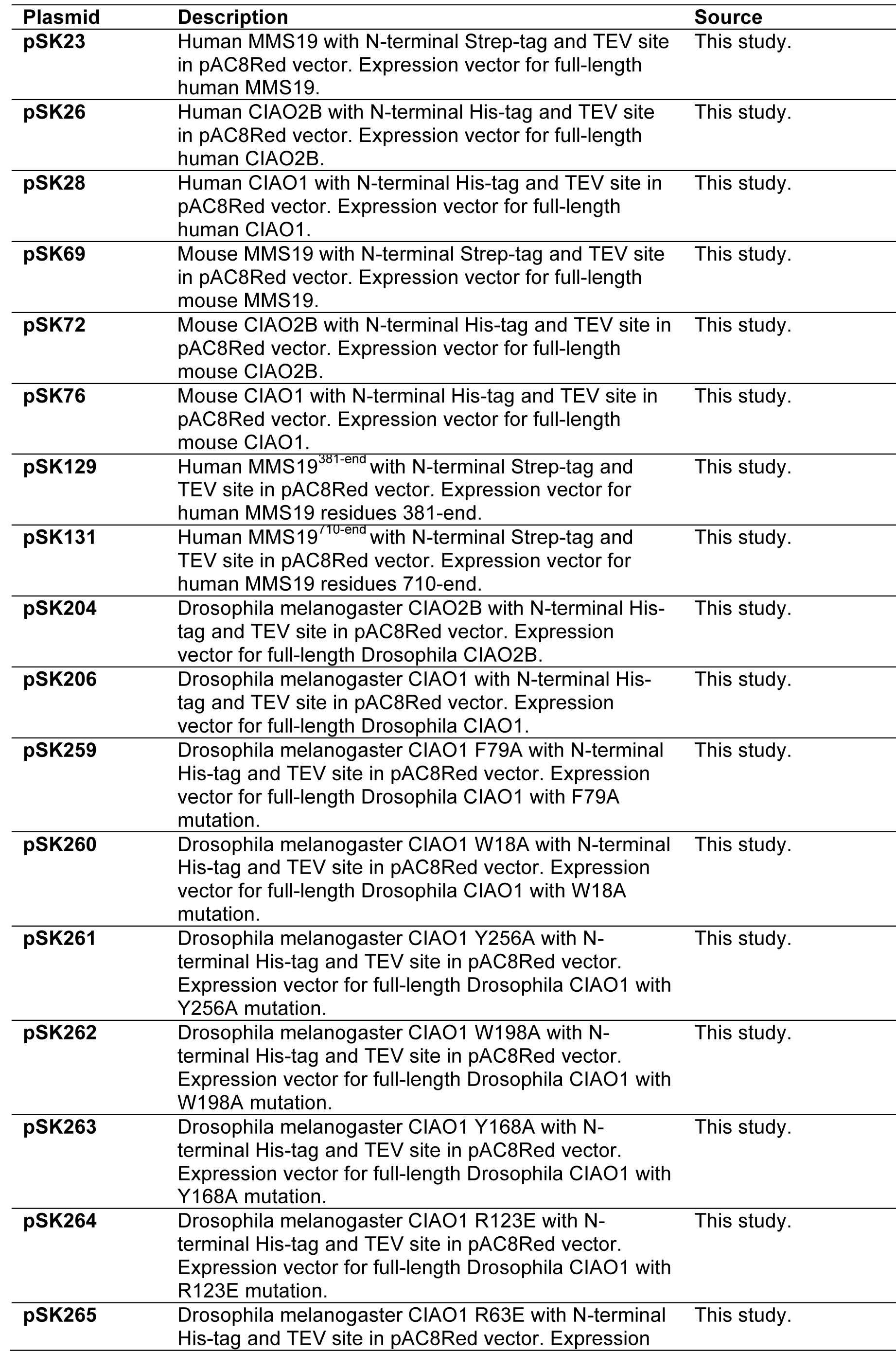
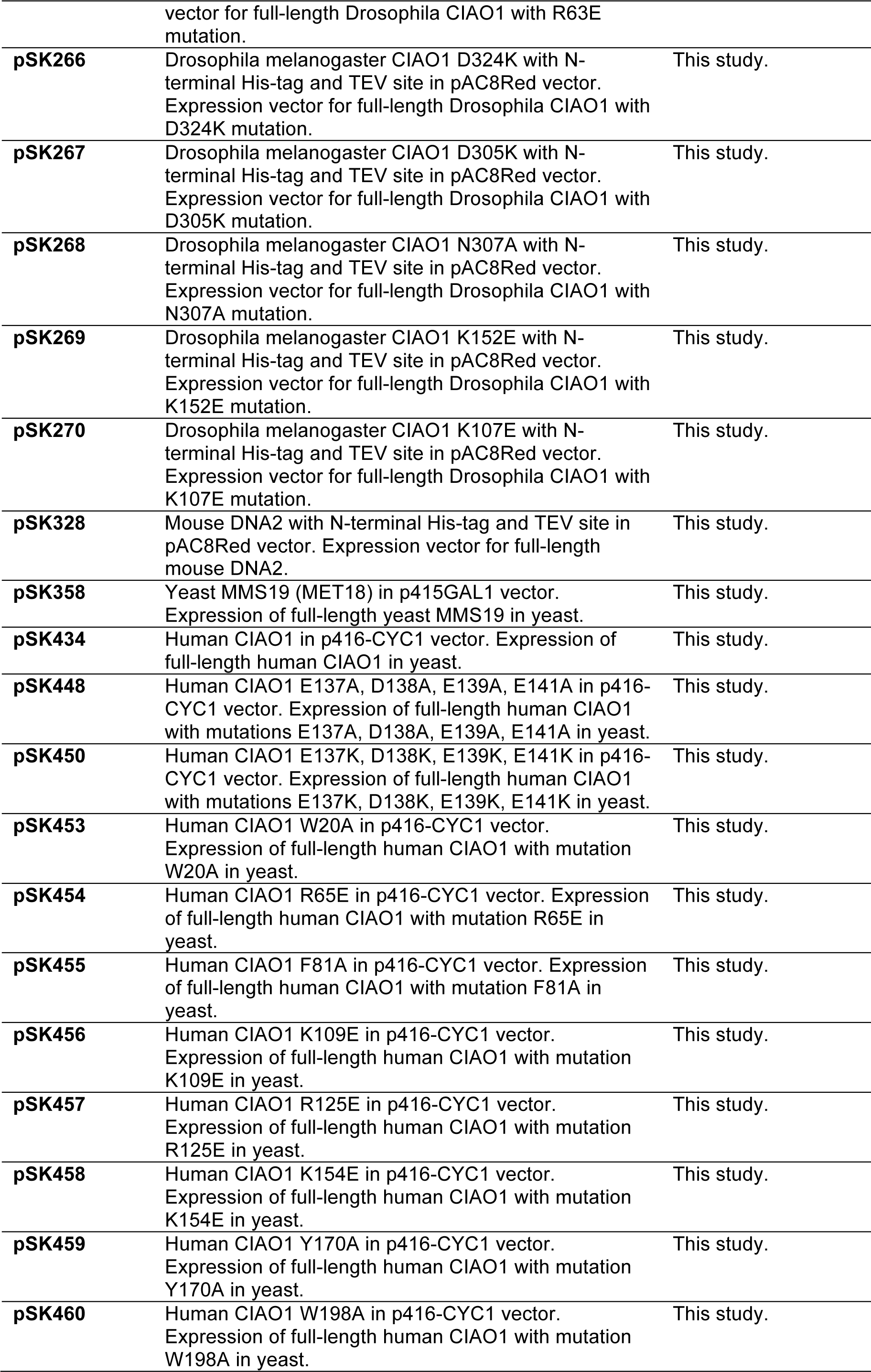
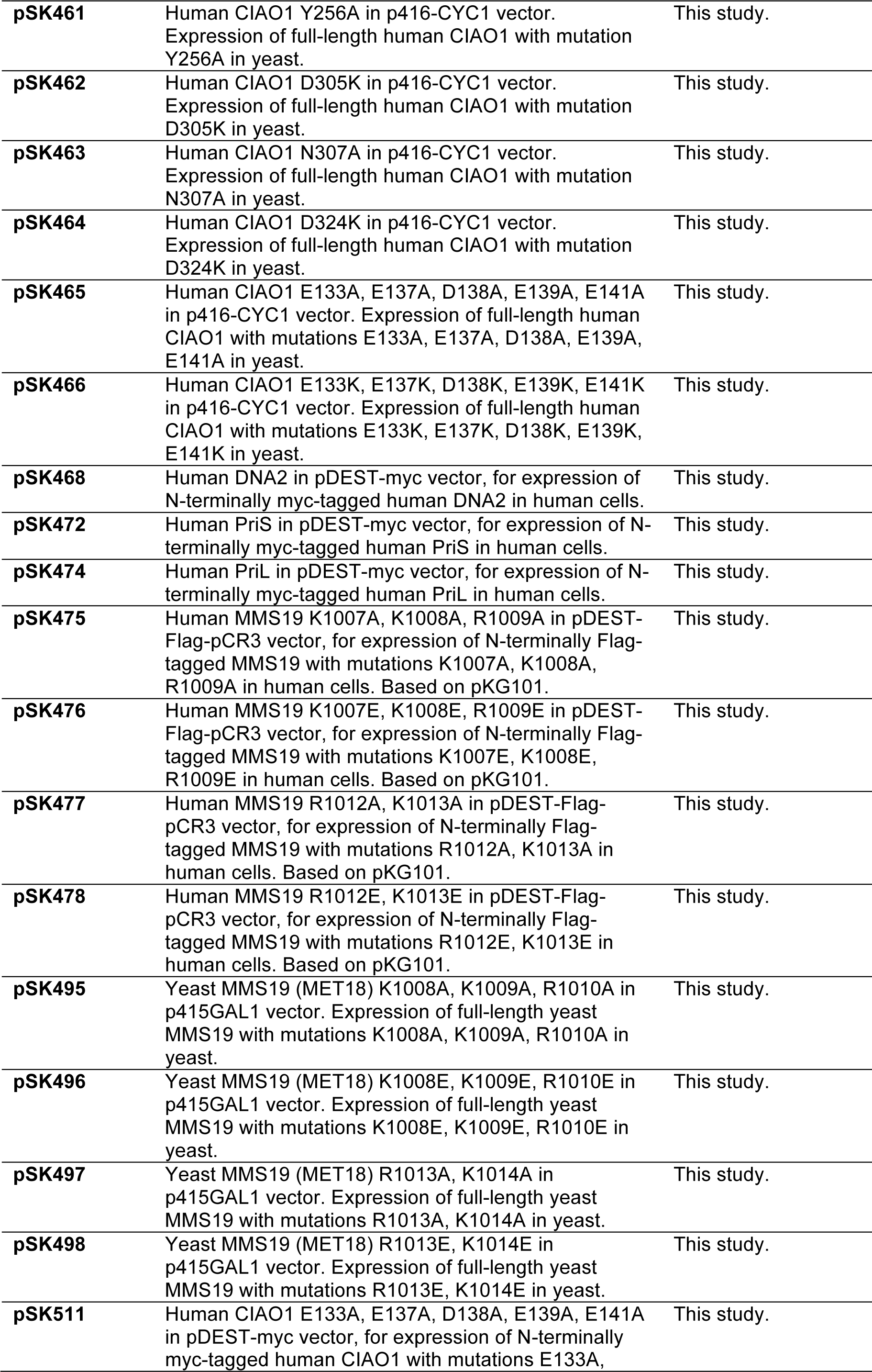
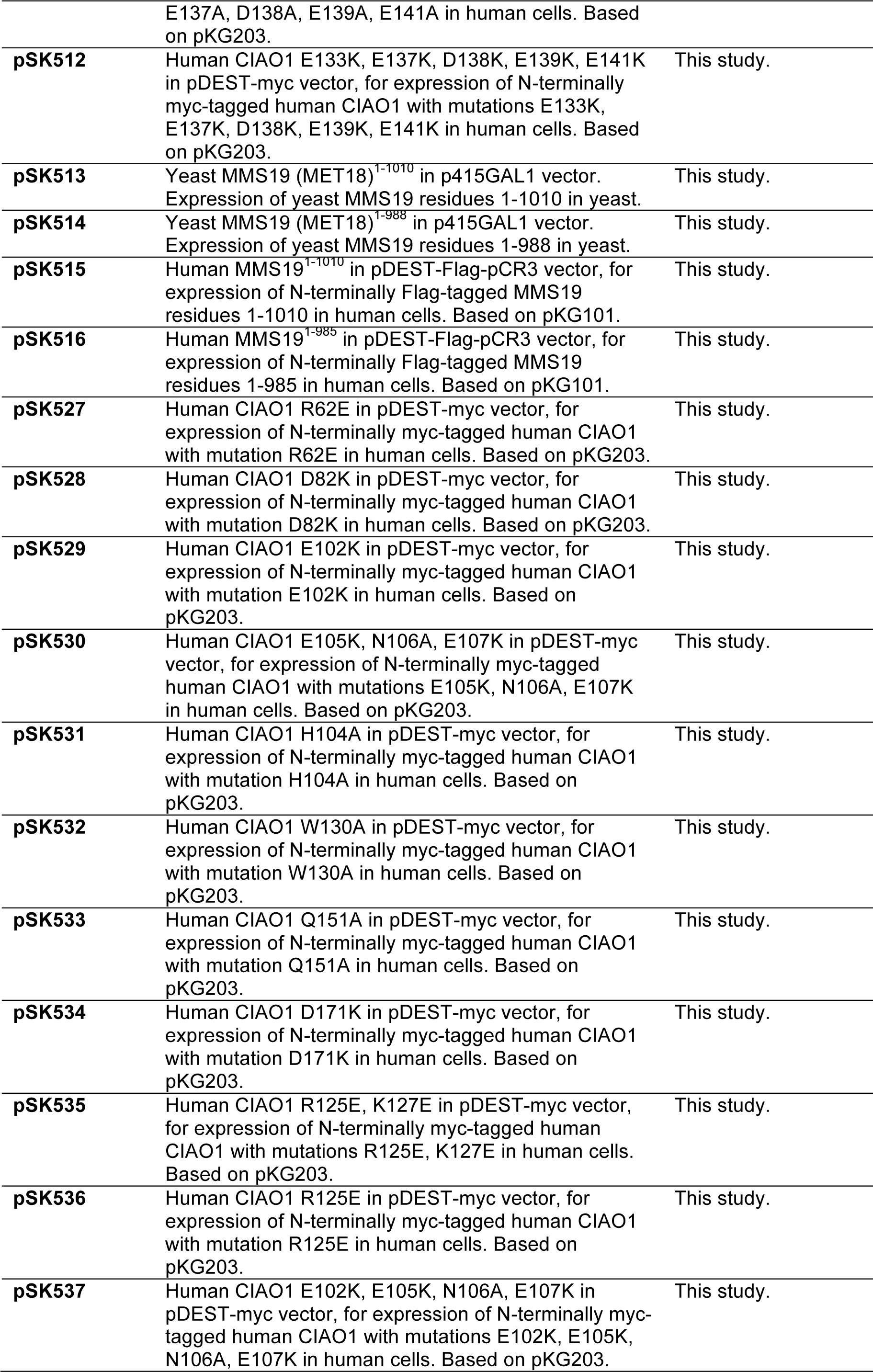
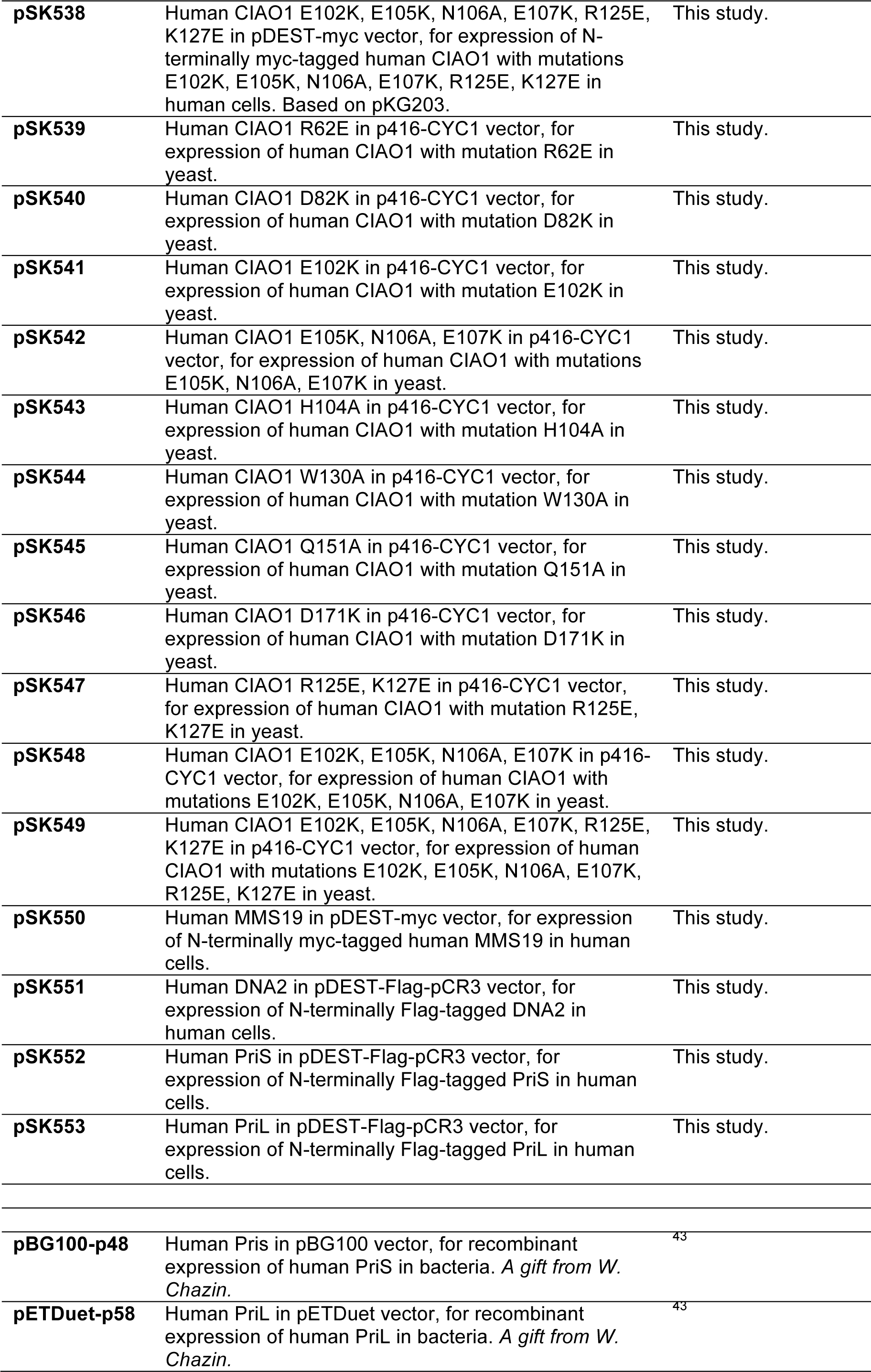
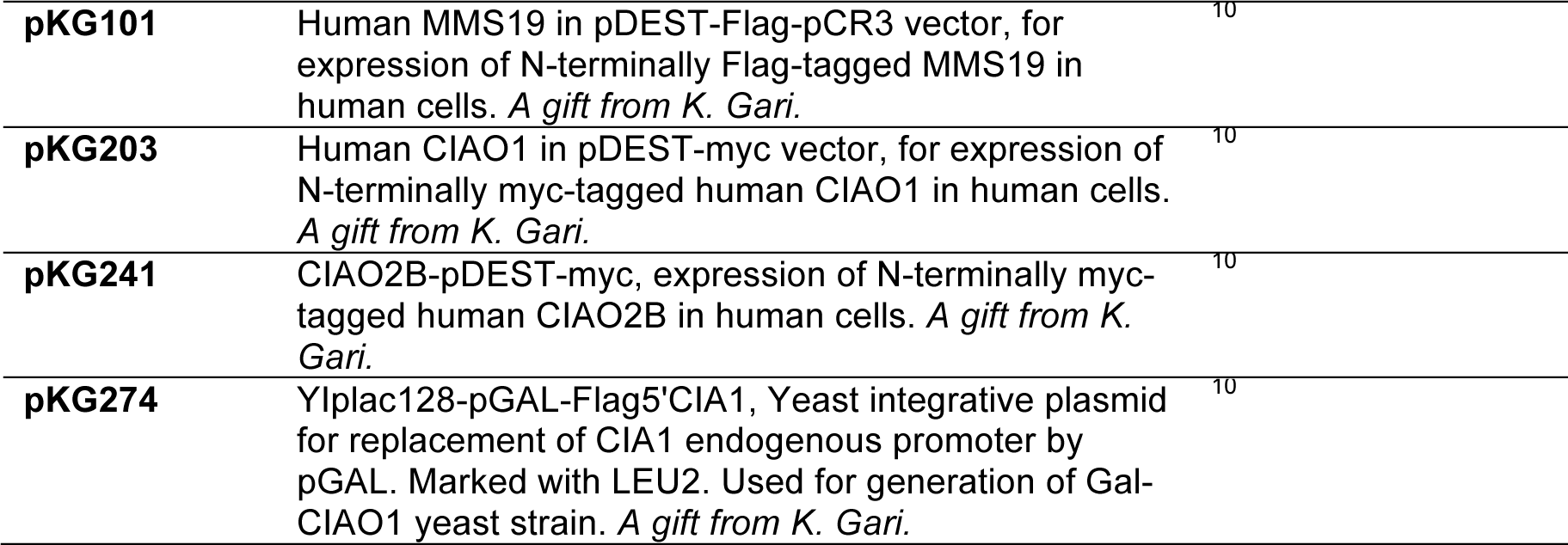
Plasmids used in this study.

**Extended Data Table 3.** Yeast strains.

| Strain | Parental strain | Genotype | Source |
| --- | --- | --- | --- |
| *Saccharomyces cerevisiae* MET18 knockout | BY4741 | MATa, his3Δ1 leu2Δ0 met15Δ0 ura3Δ0 | Dharmacon |
| *Saccharomyces cerevisiae* P<sub>GAL</sub> FlagCIA1-LEU2 | W303 | MATa, RAD5 leu2-3,112 trp1-1 can1-100 ura3-1 ade2-1 his3-11,15 | this study, based on<sup>10</sup>, using plasmid pKG274 |

